# Comparing Neuroprotective Drug Efficacy in Rodent Neonatal Brain Injury Models

**DOI:** 10.1101/2025.09.17.676833

**Authors:** John D.E. Barks, Yiqing Liu, Julie Sturza, Niko Kaciroti, William J. Meurer, Faye S. Silverstein

**Author notes:** Address for correspondence: John Barks MD, Neonatal-Perinatal Medicine, 8-621 Mott Hospital SPC 4254, University of Michigan Health System, 1540 E. Hospital Dr. Ann Arbor, MI 48109-4254.

## Abstract

**Background:** A challenge in preclinical neonatal neuroprotection research is implementation of study designs that enable direct comparison of multiple potentially effective drugs. We used adaptive design to test four FDA approved drugs (azithromycin, erythropoietin, caffeine, melatonin) concurrently and determine which best combined safety and efficacy.

**Methods:** Seven-day-old (P7) rats underwent hypoxia-ischemia (HI; right carotid ligation + timed 8% O_2_ exposure); some experiments included pre-treatment with agents that induced inflammation, and some included post-HI brief moderate hypothermia. Sensorimotor and neuropathology measures were incorporated into a Composite Score that also accounted for deaths. Outcome was initially evaluated at P21 and in confirmatory studies at P35. A pre-specified Bayesian algorithm with futility and efficacy stopping rules was devised to analyze emerging data and adjust subsequent animal allocation among drug groups.

**Results:** In all models either azithromycin or erythropoietin (EPO) offered superior neuroprotection at P21 and the other was “runner-up”. Caffeine and melatonin conferred modest neuroprotection in pure HI but were quickly eliminated in hypothermia-treated HI. At P35 azithromycin and EPO outcomes were generally similar.

**Conclusion:** These results support azithromycin or erythropoietin as candidate neuroprotective agents and warrant future studies in large animal neonatal cerebral hypoxia-ischemia models.

**Impact:**

- Our preclinical comparative efficacy strategy using adaptive design tested four clinically available drugs (azithromycin, caffeine, erythropoietin and melatonin) in multiple preclinical rodent brain injury models and consistently identified differences in efficacy among drugs.
- Three key factors differentiating our approach from existing reports were direct within-litter comparisons, evaluation in multiple injury models and inclusion of both function and neuropathology outcomes.
- Either azithromycin or erythropoietin, even with a 2-hour initiation delay, was the most neuroprotective drug in all models, with the other a close “runner-up”.
- Less effective drugs were recognized and eliminated early by the prespecified Bayesian algorithm.
- These results support azithromycin or erythropoietin as candidate neuroprotective agents and support future studies in large animal neonatal cerebral hypoxia-ischemia models.

## Introduction

Despite the advent of therapeutic hypothermia, neonatal hypoxic-ischemic encephalopathy (HIE) remains a major contributor to long-term disability^1,2^; over 40% of hypothermia-treated survivors have major neurologic deficits^3^. In high-income countries HIE is the best characterized etiology of neonatal brain injury. Although dozens of agents attenuate neonatal HI brain injury in rodent preclinical studies^4–6^, the first two drugs that were evaluated in phase 2 or 3 clinical trials (xenon, erythropoietin) did not provide incremental benefit when combined with hypothermia^3,7,8^.

These disappointing results raised a critical question for future research - how to prioritize among many available drugs for further study in larger mammals and ultimately in clinical trials. One substantial challenge stems from the understandable tendency of investigators to advocate for agents that are the focus of their laboratories. Systematic direct comparison among neuroprotective drugs in established neonatal brain injury models has been limited^9^.

A challenge that must be addressed in neuroprotection studies that directly compare efficacy among multiple drugs is implementation of experimental designs and analytic methods that acknowledge the large number of agents that could be tested. We propose a strategy to address these challenges and establish a framework to accelerate progress in neonatal neuroprotection. We selected four strong candidate drugs, based on:

i. Prioritizing FDA-approved drugs, i.e. drug repurposing, to potentially accelerate ultimate approval for neonatal neuroprotection^10^;
ii. Avoiding drugs with known safety concerns for human neonates (e.g. tetracyclines);
iii. Incorporating composite neuroprotective efficacy measures that assess survival, functional, and neuropathology outcomes;

The four drugs that met these criteria for which there is published neonatal rat dosing data were: azithromycin, erythropoietin (EPO), caffeine and melatonin. We directly compared their efficacy in rodent HI models, using a pre-specified within-litter response-adaptive randomization design with shared controls^11,12^.

Our goal with adaptive design was to evaluate multiple treatment options concurrently and determine which best combined safety and efficacy, while exposing the fewest subjects to less-safe or less-effective therapies. Pre-specified Bayesian algorithms were devised to analyze emerging trial outcome data and adjust subsequent subject allocation to the areas of greatest response or greatest uncertainty. Drugs with a low probability of superiority by Bayesian analysis could be set aside, and those with the highest probability of superiority would be strong candidates to advance to next-phase studies in larger, gyrencephalic animal models.

We used a well-characterized neonatal rat hypoxic-ischemic (HI) lesioning method (unilateral carotid ligation followed by timed exposure to moderate hypoxia) ^13^ at postnatal day 7 (P7) to model term or near-term brain injury. Use of neonatal rodents facilitates rapid sample size accrual; this model is widely used to delineate injury mechanisms^14,15^ and evaluate therapies^16,17^. In view of the potent interactions between mechanisms of hypoxic-ischemic and inflammatory neuropathology, we also evaluated inflammation-sensitized HI^18,19^. It would be very difficult to replicate 72h of therapeutic hypothermia, as is used clinically, in neonatal rats; however, brief (3h) moderate hypothermia, initiated one hour after the end of hypoxia exposure, is modestly protective^20,21^, and is included in some experiments.

## Methods

### Overview

All animal protocols were approved by the University of Michigan IACUC. Wistar rats (dams with pups) were obtained from Charles River Laboratories (Portage, MI). All animals from each litter were lesioned concurrently on postnatal day 7 (P7) and allocated to drug-treatment or saline-control groups (see *Allocation scheme* below). All animals received their initial injection of test drug or saline two hours after the end of hypoxia exposure. Since sex may influence susceptibility to brain injury and endogenous recovery mechanisms^22,23^, males and females were distributed equally between groups in each litter and for each treatment. Animals were weighed before surgery, daily for four subsequent days, and weekly beginning on P14. Rectal temperatures were measured before surgery, after hypoxia, and daily during the treatment period.

Based on results of a prior study^24^, we used a two-tiered evaluation process. Each litter included animals from each treatment group and shared controls. In the first tier, four drugs were compared together with littermate saline-injected controls, and the primary outcome was a Composite Score that incorporates survival, sensorimotor function and a brain damage severity measure at postnatal day 21 (P21) ^25^. At P21 the Composite Score relies on hemisphere weight difference as a reliable measure of ipsilateral HI tissue loss^26^ and enables rapid outcome feedback for response-adaptive randomization and efficient sample size accrual to concurrently compare multiple drugs. For each injury model, using Bayesian analysis, *a priori* efficacy criteria were applied to rapidly eliminate drugs with lower probability of being the most neuroprotective and to select the two best-performing drugs to advance to the second tier, in which the primary outcome was re-evaluated after a longer recovery (P35+). In the second tier, Composite Scores included functional and histopathology measures. Drugs and saline were labeled A-E, and individuals performing all procedures and evaluations were unaware of drug identities.

### Animal lesioning

Isoflurane-anesthetized P7 rats underwent right carotid artery ligation^27^, recovered in incubators (90 min, at 36.5°C), were placed in acrylic containers partially submerged in a water-bath (36.5°C), and exposed to a constant flow of 8% oxygen/balance nitrogen for 50, 60 or 90 min. Except for experiments that included post-HI hypothermia, they recovered in incubators (15 min, 36.5°C) and then returned to dams.

### Hypothermia protocol

For experiments that included post-HI hypothermia, after the end of hypoxia animals recovered for 1 hour in a 36.5°C incubator and then were placed in a 30°C incubator, separated by partitions for each animal to prevent huddling, for 3 h, and then returned to their dams. This protocol confers modest neuroprotection and enables detection of added effects of drug therapy plus hypothermia^28,29^. Drug injections were initiated at 1h after the start of hypothermia.

### Inflammation-amplification

Prior studies enabled us to identify practical approaches to model the impact of concurrent infection on HIE outcomes. To model concurrent Gram-negative infection and HI, P7 rats received i.p. injections of LPS (E. coli O55:B5, Millipore Sigma) 0.05 mg/kg; 150 min later they underwent right carotid artery ligation, recovered in incubators (90 min, 36.5°C), and began 50 min HI, 4h after LPS injection. To model concurrent Gram-positive infection, P7 rats received i.p. injections of the lipopeptide Pam3CysSerLys4 (PAM, Millipore Sigma) 0.5 mg/kg; 270 min later they underwent right carotid artery ligation, recovered in incubators (90 min, 36.5°C), and began 60 min HI 6h after PAM injection.

### Outcome evaluation – Functional

To detect asymmetric sensorimotor deficits, vibrissae-stimulated forepaw placing (10 trials/side, weekly from P14; full forepaw extension, 1, incomplete response 0.5, normal=10/10) and left/right ratio of forepaw grip traction strength (3 trials/side, from P21; normal score=1.0) were evaluated, as previously reported^24^. Morris Watermaze testing of spatial learning and memory^28,30,31^ was included in some second-tier studies, beginning after the completion of P35 sensorimotor testing.

Watermaze testing was conducted in the University of Michigan Animal Phenotyping Core, with a pool diameter of 120 cm, water depth of 46 cm, a 20x15 cm platform 1 cm below the water surface, using Ethovision XT video-tracking software (Noldus). Animals trained to a fixed platform location (4 trials/day, maximum duration 60 sec) over 4 days, and then a 60-sec no-platform Probe Test was performed on day 5. During Day 1-4 training trials, animals that did not find the platform within 60 sec were placed on the platform. Probe Test latency (sec) to first cross the former platform location was measured; shorter latency indicates better memory.

### Outcome evaluation – Pathology

At P21 brain damage severity was quantified by comparing bilateral hemisphere weights, which correlate closely with morphometry^26^, and % intact right hemisphere was calculated (100*right hemisphere weight/left hemisphere weight). For second-tier experiments brain damage was evaluated by quantitative morphometry (using NIH ImageJ) of cresyl violet stained coronal sections in a regularly spaced series ranging from the anterior genu to the posterior genu of the corpus callosum. Hemisphere volumes were calculated from bilateral hemisphere intact-staining areas; %Intact right hemisphere was calculated from hemisphere volumes, using the same formula as for hemisphere weights.

### Outcome evaluation – Composite Score

Rigorous preclinical neuroprotection trials should incorporate measures of survival, tissue injury, and function^32^. We applied a composite measure that incorporated these three elements, yielding a single measure that assessed survival, quantifiable measures of function, and brain damage severity^25^. To generate a 30-point Composite Score both functional outcomes and brain damage severity were each scored out of 10; normal rats consistently score 30 and animals that died after treatment but before testing were scored as 0. Scores were calculated by adding: [left forepaw vibrissae score (maximum=10)] + [(L/R forepaw grip ratio) x 10] + (%Intact right hemisphere / 10).

### Candidate Drugs

We utilized within-litter randomization to minimize the impact of between-litter variability in lesion severity^21^. First-tier experiments included four drugs; this decision was based on: (i) typical litter size is 10-12 rats, (ii) desire to assign at least one male and one female to each treatment group in each litter, (iii) need for shared controls of both sexes in each litter. We selected four drugs that are already used in human neonates and for which neonatal rodent dosing, including repeated-dose regimens, has been published: azithromycin, caffeine, erythropoietin and melatonin. Regimens and references are presented in Table 1. Drug solutions were prepared by a member of another laboratory, and labels were coded A-E. The first injection of drug or saline was at two hours after the end of hypoxia exposure; in hypothermia groups the first injection was after 1h of hypothermia. This delayed onset of treatment would likely be necessary in a clinical trial to evaluate any of these drugs. Drugs and saline were administered by intraperitoneal (i.p.) injection at 2, 24, 48, 72 and 96 hours after the end of HI.

**Table 1:** Drug Regimens.

| **Drug** | **Citations** | **Presumptive Mechanism(s)** | **Proposed Regimen** | **Clinical <sup>a</sup>** |
| --- | --- | --- | --- | --- |
| Azithromycin | 24,25 | Immunomodulation | 45 mg/kg 2h, 22.5 mg/kg 24, 48, 72, 96h | +/- |
| Caffeine citrate | 33-35 | Adenosine receptor antagonist, antioxidant | 20 mg/kg 2, 24, 48, 72, 96h | ++ |
| Erythropoietin | 36-41 | Neurogenesis, angiogenesis, oligodendrogenesis | 5000 u/kg 2, 24, 48, 72, 96h | ++ |
| Melatonin | 42,43 | Antioxidant, anti-inflammatory, anti-apoptotic | 15 mg/kg 2, 24, 48, 72, 96h | + |

a. The “Clinical” column estimates frequency of drug use (for any indication) in clinical neonatology (++: common; +: selected patients; +/-: occasional).

### Allocation scheme

Cohorts of 1 or 2 litters (n=11-12/litter) were concurrently lesioned. We used Fixed and Adaptive Clinical Trial Simulator (FACTS, Berry Consultants, Austin, TX) to design Tier 1 with four drug arms and a shared control arm (see Supplement 1). P21 outcomes in Tier 1 were selected to provide timely, pragmatic, reliable initial results. Tier 1 allocation is illustrated in **Table 2**. Drug identification (i.e. un-masking to treatment identity) only occurred after the conclusion of statistical analyses of all outcomes.

**Table 2:** Tier 1 Drug Group Allocations.

| Injury Model | Drug | Animals/Group/Cohort <sup>a</sup> |  |  |  |  | Total |
| --- | --- | --- | --- | --- | --- | --- | --- |
|  |  | Cohort 1 | Cohort 2 | Cohort 3 | Cohort 4 | Cohort 5 |  |
| HI90 |  |  |  |  |  |  |  |
|  | A | 5 | 5 | 2 |  |  | 12 |
|  | B | 5 | 5 | 2 |  |  | 12 |
|  | C | 4 | 5 | 2 |  |  | 11 |
|  | D | 5 | 4 | 3 |  |  | 12 |
|  | E | 4 | 5 | 3 |  |  | 12 |
| HI90+HT |  |  |  |  |  |  |  |
|  | A | 5 | 3 | 0 | 0 | 0 | 8 |
|  | B | 5 | 12 | 13 | 12 | 18 | 60 |
|  | C | 5 | 6 | 6 | 6 | 6 | 29 |
|  | D | 5 | 0 | 0 | 0 | 0 | 5 |
|  | E | 4 | 3 | 5 | 6 | 0 | 18 |
| LPS+HI50 |  |  |  |  |  |  |  |
|  | A | 2 | 5 | 0 |  |  | 7 |
|  | B | 2 | 5 | 3 |  |  | 10 |
|  | C | 3 | 4 | 6 |  |  | 13 |
|  | D | 3 | 4 | 0 |  |  | 7 |
|  | E | 2 | 6 | 15 |  |  | 23 |
| PAM+HI60 |  |  |  |  |  |  |  |
|  | A | 4 | 0 |  |  |  | 4 |
|  | B | 5 | 0 |  |  |  | 5 |
|  | C | 5 | 6 |  |  |  | 11 |
|  | D | 5 | 0 |  |  |  | 5 |
|  | E | 5 | 18 |  |  |  | 23 |
| LPS+HI50+HT |  |  |  |  |  |  |  |
|  | A | 5 | 0 | 0 |  |  | 5 |
|  | B | 5 | 5 | 3 |  |  | 13 |
|  | C | 4 | 6 | 6 |  |  | 16 |
|  | D | 5 | 0 | 0 |  |  | 5 |
|  | E | 5 | 9 | 15 |  |  | 29 |
| PAM+HI60+HT |  |  |  |  |  |  |  |
|  | A | 4 | 0 |  |  |  | 4 |
|  | B | 5 | 10 |  |  |  | 15 |
|  | C | 5 | 6 |  |  |  | 11 |
|  | D | 5 | 0 |  |  |  | 5 |
|  | E | 5 | 8 |  |  |  | 13 |

a. Shaded cells represent cohorts with response-adaptive allocation

### Statistical Methods

We evaluated six lesioning protocols in first-tier experiments: HI alone (N=59; 5 litters), HI plus hypothermia (HT) (N=120; 10 litters), LPS-amplified HI (N=60; 5 litters), PAM-amplified HI (N=48; 4 litters), LPS-amplified HI plus hypothermia (N=68; 6 litters), PAM-amplified HI plus hypothermia (N=48; 4 litters). We defined composite outcome scores as: “Very Effective”, 26 ±3 (mean ±SD), “Moderately Effective”, 17 ±4, “Untreated control”, 11 ±4, and applied a design similar to adaptive clinical trials ^11,44,45^, with pre-specified interim looks after each cohort, and pre-specified “win” and “futility” criteria (see **Supplement 1, Design and Simulation Report**). Based on Composite Scores at each interim look, the algorithm first decided if a drug met criteria of ≥2.3 SD above control scores. Derived in numerical simulations, this threshold balanced accurate identification of superior drugs with not advancing drugs with lower effect size. Next, it decided if “futility” criteria were met (i.e. despite more cohorts, no drug would be declared maximally effective vs. the other drugs). If neither was true, for the next cohort the algorithm applied response adaptive animal allocation to favor drugs with better chances of superiority. If the algorithm-generated allocation for a drug was <0.1 (i.e. 10% of a cohort), it was eliminated from that cohort onward. The maximum first-tier sample size was set at 120, and we recognized that the high neuroprotective efficacy of two or more drugs might not be distinguishable. Tier 1 outputs for each drug were the probabilities that it was superior to control (P_sup_; threshold 0.99) and that it was the most neuroprotective drug (P_max_; threshold 0.95).

In a secondary, frequentist, analysis to evaluate for differences in relative neuroprotective efficacy among the 4 drugs, we used either ANOVA, or a mixed model controlling for clustering within litters. Additionally, all experiments with normothermic recovery were pooled and then compared to the pooled experiments that included treatment with hypothermia, using a mixed model controlling for clustering within litters, where drug (A, B, C, D, E) predicts composite score, controlling for injury model (e.g. HI90 vs. LSP+HI50 vs. PAM+HI60). Post-hoc pairwise comparisons were conducted with Tukey-Kramer multiple comparisons adjustment.

In all experiments, mortality was compared among groups by Fisher’s exact test. The effects of drugs on serial temperatures or weights were compared among groups using mixed-effects models (SAS Proc Mixed, SAS, Cary NC) with rats clustered within litters. Post-hoc pair-wise comparisons were done to determine differences between drug groups and controls, using the Tukey-Kramer multiple comparison adjustment.

In second-tier experiments, that began with animals that underwent HI alone (N=36), followed by HT-treated HI (N=48) inflammation-amplified HI (N=36) and HT-treated inflammation-amplified HI (N=24), we advanced the drug with the leading probability of maximal efficacy (P_max_) at P21, and the first “runner-up”, for each injury model. We used first-tier effect sizes and SD values to estimate second-tier sample sizes needed to determine if at least one drug was superior to controls (8-16/group). Group differences were evaluated using frequentist statistical methods, with t-test, ANOVA, or nonparametric tests, as appropriate (Prism, GraphPad Software, Boston, MA).

## Results

### Physiology measures

Mortality was very low with hypoxia-ischemia alone, but higher with LPS-induced inflammation-amplification, as detailed below. Weight gain trajectories in the drug groups did not differ from controls for any injury condition (see Supplementary Data Table 1). Serial temperatures in the drug treatment groups did not differ from controls for any injury condition (Supplementary Data Table 2). Sex had no impact on any outcome, under any lesioning conditions.

### Tier 1 outcomes

#### Hypoxia-Ischemia

The first experiment, in which pups were allocated equally to five groups (N=23), revealed that Drug E (azithromycin) had a Bayesian probability of maximal efficacy (P_max_) of 0.996; this value exceeded the *a priori* success threshold of 0.95, and according to our Bayesian protocol no additional experiments were needed. However, we added three more litters with equal group allocation (final n=59) to confirm this result using frequentist methods.

Composite Scores for all four drugs were superior to controls, and azithromycin remained superior to the other drugs (p<0.0001 ANOVA with post-hoc Šidàk’s multiple comparisons test; see Fig. 1 d). We also evaluated the three components of the composite score separately.

**Figure 1:**
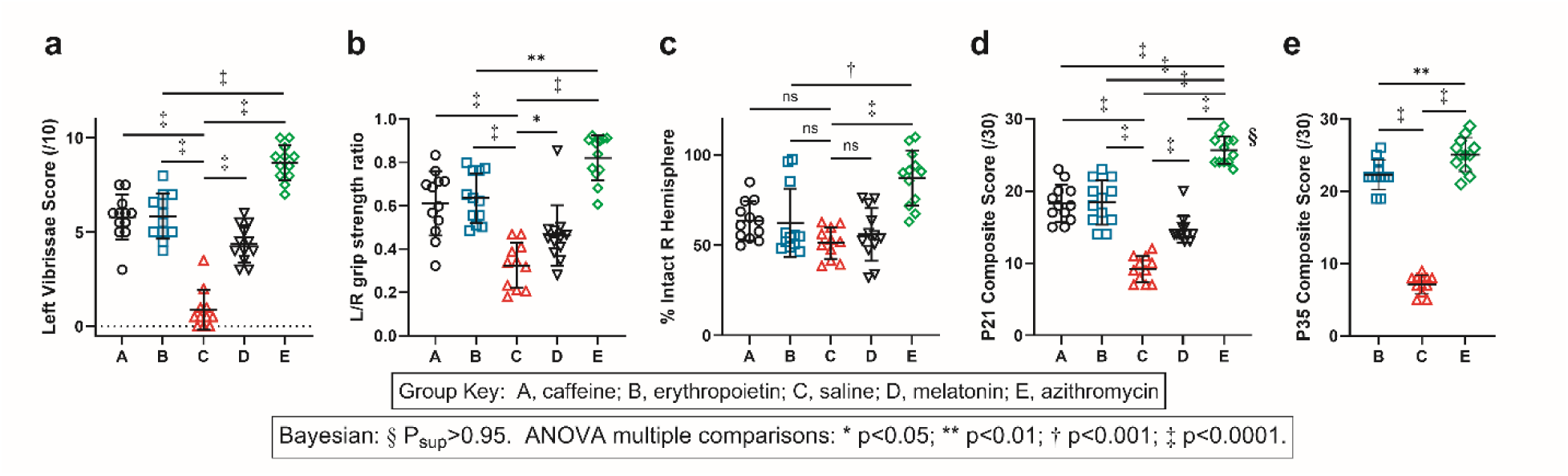
Comparative Neuroprotective Efficacy of Four Clinically Available Drugs in Neonatal Rat Hypoxic-Ischemic (HI) Brain Injury Model: For the first two experiments, animals were randomly allocated (4-5/group) into four drug treatment groups or shared saline controls. After right carotid artery ligation, postnatal day 7 (P7) rats spent 90 min in chambers with 8% O_2_ (HI90). Once daily injections over 5 consecutive days began 2h after HI (see Table 1 for drug doses). Outcome evaluators were unaware of treatment identity until the end of the study. Forepaw sensorimotor function (a, b) and %intact right hemisphere tissue (c) were evaluated on P21, and composite scores (maximum 30/30) were calculated (d). Based on composite scores, after the first two experiments (N=23) the Bayesian probability of superiority (P_max_) for Drug E was 0.996, meeting *a priori* success criteria (d §, P_max_>0.95). We added three litters with equal random allocation to confirm the result using frequentist methods (final N=11-12/group); Drug E mean composite scores were higher than for the other 3 drugs (‡p<0.0001, ANOVA with post-hoc Šidàk’s multiple comparisons test), but scores for all drugs were higher than controls (‡p<0.0001, ANOVA post-hoc test). Based on the 2 functional outcomes, all drugs improved outcome relative to controls (a, b), and Drug E was superior to drug B (‡ p<0.0001, ** p<0.005, Šidàk’s post-test). Based on neuropathology scores (c), only Drug E was neuroprotective (‡ p<0.0001 vs. controls, ANOVA post-test), and Drug E was also superior to drug B (†p<0.001 ANOVA with post-hoc test). We then evaluated juvenile-age outcomes on P35 (e) in rats that were randomized on P7 to Drug B, Drug E or saline (C) (12/group); in addition to sensorimotor testing, %intact right hemisphere tissue by volume was calculated (see Methods). At P35 composite scores in Drug E-treated animals were higher than with Drug B (**p<0.005, ANOVA with Tukey’s post-test). (bars: mean ±SD). Drug Group key: A, caffeine; B, erythropoietin; C, saline; D, melatonin; E, azithromycin.

Vibrissae scores for all 4 drugs were superior to controls, and azithromycin performed best (p<0.005 ANOVA with post-hoc Šidàk’s test; Fig. 1 a, b); based solely on evaluation of tissue damage only azithromycin was superior to controls (Fig. 1 c).

#### HI plus Hypothermia

After evaluating Composite Scores from the first two litters (N=24), the Bayesian algorithm dropped the Drug D regimen (melatonin). Animals in 8 additional litters were treated with drugs allocated by response-adaptive randomization, based on Composite Scores in preceding litters; the *a priori* superiority threshold was crossed at 120 animals and Drug B (EPO) was declared the winner. Drug A (caffeine) was eliminated after the fourth litter. Final group sizes were 8, 60, 29, 5 and 18 respectively for caffeine, EPO, control, melatonin, and azithromycin. The mortality rates were 1/8, 0/60, 3/29, 0/5 and 2/18 respectively, and only Drug B (EPO) mortality rates were lower than controls (p<0.05, Fisher Exact test). The P_max_ for EPO was 0.9826; by *post-hoc* frequentist analysis, the least squares mean Composite Scores for both EPO and azithromycin were higher than controls (p<0.0001 Mixed Model, with post-hoc Tukey- Kramer adjustment; Fig. 2 d), and the scores for EPO and azithromycin did not differ (p=0.62, Tukey-Kramer; Fig. 2 d). A summary comparison of Bayesian and Frequentist analysis results in all Tier 1 experiments is presented in **Table 3**.

**Figure 2:**
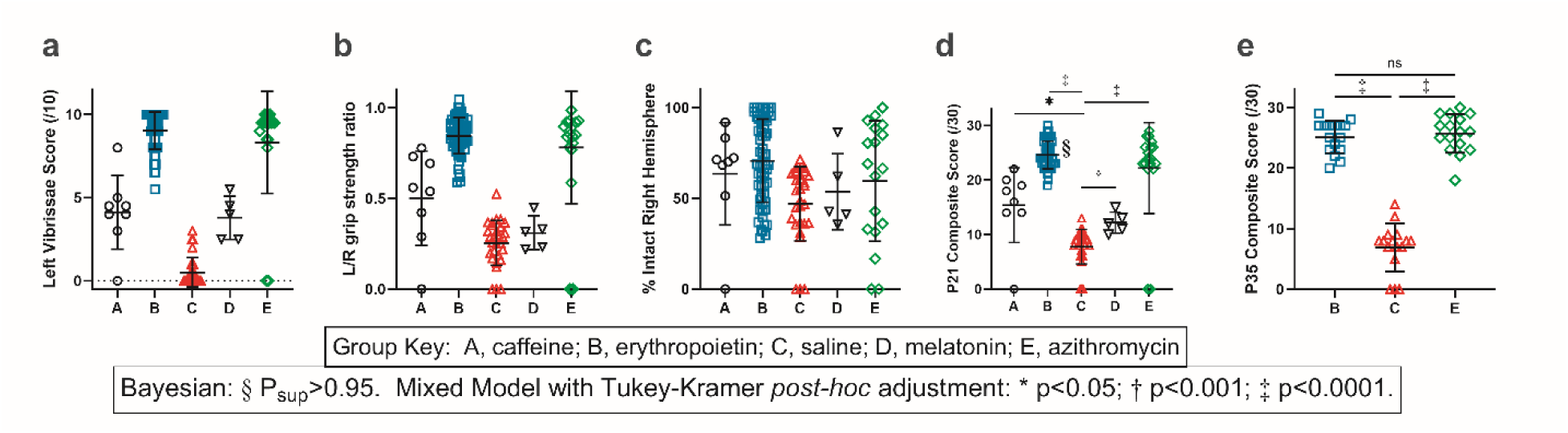
Adding post-HI brief hypothermia modifies the neuroprotective efficacy of 4 clinically available drugs in neonatal rat brain hypoxia-ischemia (HI) model: After right carotid artery ligation, postnatal day 7 (P7) rats spent 90 min in chambers with 8% O_2_ (HI90), followed 1h later by 3h hypothermia (HT) in a 30 °C incubator. Regimens of one injection/day over 5 consecutive days began 2h after HI, with the initial litters (N=24) evenly randomized to one of 4 drug groups (A, B, D, E) or controls (C). In Fig. 2 a-d outcomes on P21 were evaluated as in Fig. 1 a-d. Based on the Bayesian algorithm Drug D was dropped after the two initial even-allocation litters, and after two additional litters allocated by response-adaptive randomization (N=24) Drug A was dropped. In 3 subsequent pairs of litters group allocation (to B, C or E) was by response-adaptive allocation. After 120 animals the algorithm declared Drug B (erythropoietin) the “winner” (**d**, §: P_max_ 0.9826). By *post-hoc* frequentist analysis, least squares mean composite scores for B and E did not differ (p=0.62, Tukey-Kramer) (bars: mean ±SD). The Bayesian algorithm, as specified, readily eliminated less-effective drugs (A, D). However, under conditions in which we modeled the clinical scenario that included TH after HI, it was much more challenging to declare one clear winner between two drugs with very similar efficacy. We also evaluated P35 outcomes (**e**). 48 P7 rats underwent HI90 and were evenly randomized (within-litter) to 2 drug groups, (B, E) or controls (C). After functional testing, animals were sacrificed; % intact right hemisphere tissue was calculated using hemisphere volumes (see Methods) and composite scores were calculated. The neuroprotective efficacy of both B and E relative to controls was sustained (**e**, **‡**p<0.0001, ANOVA post-hoc test), but there was no difference in mean composite score between the two drug groups. Cognitive testing (not shown) did not alter outcomes. Drug Group key: A, caffeine; B, erythropoietin; C, saline; D, melatonin; E, azithromycin.

**Table 3:** Comparison of Bayesian and Frequentist Assessments of P21 Outcomes.

| Condition | Bayesian Analysis |  |  |  | Frequentist Analysis |  |  |  |  |  |  |  |  |  |
| --- | --- | --- | --- | --- | --- | --- | --- | --- | --- | --- | --- | --- | --- | --- |
|  | Winner | P<sub>max</sub> | N<sup>a</sup> | Median Composite Score, Winner | overall P value, mixed model<sup>b</sup> | Least Squares Means Composite Scores, by condition and drug p value<sup>c</sup>, Tukey-Kramer *post-hoc* vs. NS controls (C) |  |  |  |  |  |  |  |  |
|  |  |  |  |  |  | C: NS control | CAF | p<sup>c</sup> | EPO | p<sup>c</sup> | MEL | p<sup>c</sup> | AZ | p<sup>c</sup> |
| HI90 | AZ | 0.99 | 23 | 25.5 | <0.0001 | 9.8 | 15.7 | <0.0001 | 22.5 | <0.0001 | 14.3 | 0.0005 | 26.8 | <0.0001 |
| LPS+HI | AZ | 0.97 | 60 | 28 | <0.0001 | 9.8 | 14.7 | 0.3346 | 23.9 | 0.0002 | 13.7 | 0.5463 | 27.2 | <0.0001 |
| PAM+HI | AZ | 0.97 | 48 | 27 | <0.0001 | 11.2 | 18.6 | 0.0003 | 25.4 | <0.0001 | 18.1 | 0.0016 | 27.3 | <0.0001 |
| **Normothermia post-HI, pooled** |  |  | **131** |  | **<0.0001** | **10.0** | **17.8** | **<0.0001** | **22.0** | **<0.0001** | **15.4** | **<0.0001** | **26.7** | **<0.0001** |
| HI90+HT | EPO | 0.98 | 120 | 24 | <0.0001 | 7.8 | 15.2 | 0.0021 | 24.4 | <0.0001 | 12.8 | 0.1507 | 22.8 | <0.0001 |
| LPS+HI+HT | AZ | 0.96 | 68 | 27 | <0.0001 | 9.5 | 6.8 | 0.8499 | 23.9 | <0.0001 | 9.1 | 0.9999 | 27.1 | <0.0001 |
| PAM+HI+HT | EPO | 0.88 | 48 | 28 | <0.0001 | 12.7 | 20.2 | <0.0001 | 26.5 | <0.0001 | 19.0 | <0.0001 | 25.4 | <0.0001 |
| **Hypothermia treatment post-HI, pooled** |  |  | **236** |  | **<0.0001** | **9.7** | **14.4** | **0.002** | **25.5** | **<0.0001** | **13.7** | **0.02** | **25.6** | **<0.0001** |

Drug Group key: Caffeine (CAF, drug A); Erythropoietin (EPO, drug B); Saline controls (C); Melatonin (MEL, drug D); Azithromycin (AZ, drug E).
a. N: Total number of animals at which Bayesian analysis was completed. For HI90 36 animals were added with even allocation (total N=59) for validation with frequentist analysis.
b. Linear Mixed Model P-value (a planned secondary outcome) comparing among 5 groups (Drugs A, B, D, E and shared Controls)
c. Linear Mixed Model with Tukey-Kramer multiple comparisons test, in which each of the 4 drug groups is compared with saline-injected controls (C); p-value is shown for all *post-hoc* pairwise comparisons.

#### Inflammation-amplified HI

In the LPS-amplified HI model, in one of the initial litters there were multiple early pup deaths; data from this litter was excluded, and two more even-allocation litters were added. Based on data from three litters, the algorithm dropped drugs A and D, and allocated 15 rats to Drug E, 3 to B, and 6 to controls for the next two litters. The P_max_ for azithromycin was 0.9672 (see Fig. 3 a), meeting success criteria; final allocations were 7, 10, 13, 7, and 23 for A, B, C, D and E respectively. At least one death (range 1-2) was noted in each group, except for the azithromycin group, with no deaths. By *post-hoc* frequentist analysis, mean Composite Score was higher than control for both azithromycin and EPO (p<0.001, p<0.0001, respectively, Mixed Model with Tukey-Kramer adjustment), the mean Composite Score for azithromycin was higher than for caffeine or melatonin (p<0.0001, Šidàk’s test), and azithromycin and EPO Composite Scores were similar (see Fig. 3 a).

**Figure 3:**
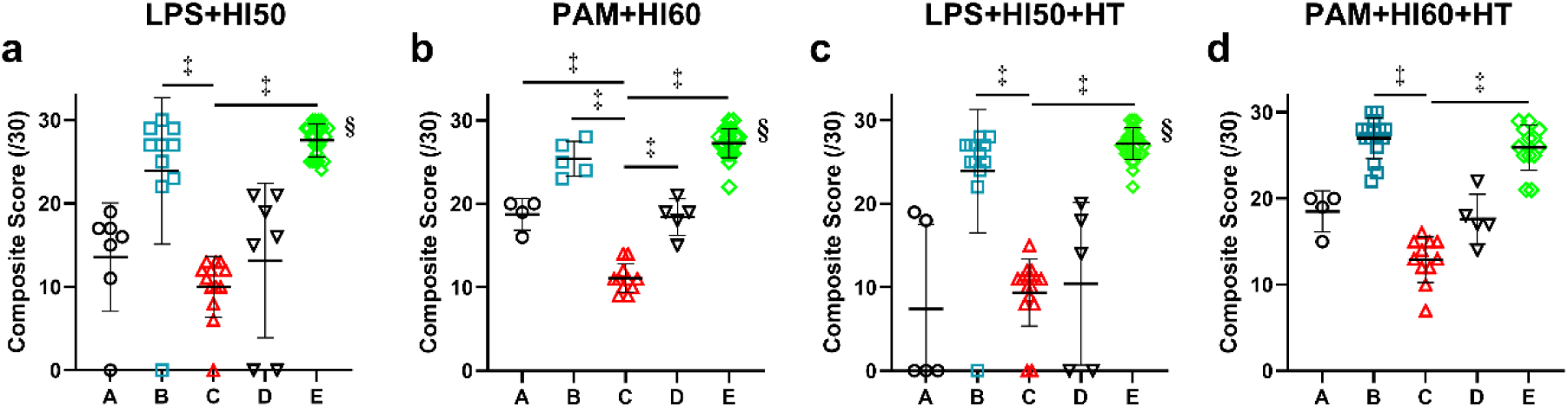
Inflammation-amplified Hypoxia-Ischemia (HI); P21 Outcomes: P7 rats received i.p. injections of LPS 0.05 mg/kg (N=60) 4 h before 50 min HI (**a**), or PAM 0.5 mg/kg (N=48) 6 h before 60 min HI (**b**), and recovered in room air. Additional animals received LPS and 50 min HI (N=68, **c**), or PAM and 60 min HI (N=48, **d**), and then 1 h after HI, underwent hypothermia (HT) for 3h in a 30°C incubator. Five-day regimens of one injection/day of Drugs A, B, D or E, or saline (C) were initiated at 2 h after the end of HI. Outcomes on P21 were evaluated as in Fig. 1. In animals that underwent LPS-amplified HI (**a**), Drugs A and D were eliminated after the initial even-allocation litters, and the algorithm stopped allocation for superiority of Drug E after 5 litters (60 animals total; §P_max_ = 0.9672). In groups A-D there was at least one death (range 1-2); there were no deaths in group E. By *post-hoc* frequentist analysis both Drugs B and E had higher mean Composite Scores than controls (†p<0.001, ‡p<0.0001 respectively, Mixed Model with Tukey-Kramer adjustment), but their scores were not statistically different. In the PAM-amplified HI experiments (**b**) Drugs A, B and D were dropped by the algorithm after the initial even-allocation litters, and allocation stopped after 4 litters (48 animals) for superiority of Drug E (§P_max_ = 0.9676). By frequentist *post-hoc* analysis, mean P21 composite scores for all 4 drugs were higher than controls (**p<0.01, ‡p<0.0001, Tukey-Kramer), and there was no statistical difference between B and E. With LPS+HI followed by HT (**c**), Drugs A and D were eliminated after the initial two even-allocation litters, and the algorithm stopped allocating after 6 litters (69 animals) for superiority of Drug E (§P_max_ = 0.9594). By *post-hoc* frequentist analysis both Drugs B and E had higher mean composite scores than controls (**c.** ‡p <0.0001, Mixed Model with Tukey-Kramer adjustment) and their scores were not statistically different. With PAM+HI followed by HT (**d**), Drugs A and D were eliminated after the initial 2 even-allocation litters, and the algorithm allocated 10, 6 and 8 animals to B, C and E respectively in the next 2 litters. After 4 litters (N=48) drug B (erythropoietin) had a P_max_ of 0.88, but had only a 1-point advantage in mean composite score over drug E (azithromycin). Although the P_max_ threshold of 0.95 had not been reached, based on our experience in the HI + hypothermia experiments (N=120), there was little scientific rationale to expend additional animals. By *post-hoc* frequentist analysis both Drugs B and E had higher mean composite scores than controls (‡p<0.0001, Mixed Model, Tukey-Kramer adjustment), but their scores were not statistically different. Drug Group key: A, caffeine; B, erythropoietin; C, saline; D, melatonin; E, azithromycin.

In the PAM-amplified HI model, after evaluation of two even-allocation litters, the algorithm allocated 18 rats to azithromycin and 6 to controls; the other drugs were eliminated. Final group allocations were 4, 5, 11, 5 and 23 to A, B, C, D and E, respectively and there were no deaths.

Drug E (azithromycin) met success criteria with a P_max_ of 0.9676 (see Fig. 3 b). By *post-hoc* frequentist analysis, the mean Composite Score for each drug was higher than controls, and the mean Composite Score for Azithromycin was higher than for caffeine or melatonin (p<0.0001 Mixed Model, with p<0.001, Tukey-Kramer) while azithromycin and EPO Composite Scores were similar (Fig. 3 b).

In prior studies in rats and piglets, hypothermia did not confer benefit vs LPS-amplified HI^46,47^ but additive benefit was described in PAM-amplified HI^48^. We evaluated the four study drugs in animals that underwent LPS-amplified HI plus delayed-initiation hypothermia. After the initial pair of even-allocation litters, the algorithm eliminated caffeine and melatonin (Drugs A and D). The algorithm stopped allocation after a total of 6 litters when azithromycin (Drug E) met superiority criteria (P_max_ = 0.9594). The final pup allocations were 5, 13, 16, 5, and 29 to regimens A, B, C, D and E, respectively. By *post-hoc* frequentist analysis, least squares mean Composite Scores for azithromycin and EPO were not statistically different. Mortality was 3/5, 1/13, 2/16, 2/5 and 0/29 in regimens A, B, C, D and E, respectively; with lower mortality in the azithromycin group than in the caffeine (A) or melatonin (D) groups (p<0.005 and p<0.05 respectively, Fisher’s exact test). In separate experiments, animals underwent PAM-amplified HI on P7, plus delayed-initiation hypothermia (Fig. 3 d). After the initial pair of even-allocation litters, the algorithm eliminated groups A and D, and allocated 10, 6 and 8 animals to B, C and E, respectively. After 4 litters the P_max_ for erythropoietin was 0.8758 but the least squares mean Composite Score for erythropoietin (26.5) was only slightly higher than for azithromycin (25.4). We judged that there was little scientific value in testing more animals when these two treatments were of similar efficacy, and neither crossed the *a priori* superiority threshold.

In a secondary analysis, using mixed models to control for clustering within litters, we evaluated relative efficacy of the 4 drugs for animals with normothermic recovery and separately in those treated with post HI hypothermia, using all three injury models. Among models with normothermic recovery, all drugs were superior to saline. Azithromycin was more effective than the other three drugs, controlling for model, EPO was superior to caffeine and melatonin and the caffeine and melatonin regimens were of similar efficacy. Among models with hypothermia treatment, all drugs were superior to saline, but EPO was superior to the other 3 drugs. Least squares mean estimates for the Composite Score for each group, which account for clustering of animals within litters and differential sample sizes among groups, are shown in **Table 3**.

### Tier 2 Outcomes

In tier one experiments, EPO and azithromycin yielded similar benefits overall, and tier 2 studies included these two drugs; outcomes were evaluated using frequentist methods.

#### Hypoxia-Ischemia (HI)

Animals underwent 90 minutes HI with normothermic recovery on P7, and received 5-day regimens of EPO, azithromycin or saline (12/group) beginning 2h after HI. Sensorimotor and brain damage outcomes were evaluated on P35. Composite Scores on P35 were higher in azithromycin-treated than in EPO-treated animals (mean ±SD: azithromycin 25 ±2, EPO 22 ±2, p=0.006, t-test), with both groups performing better than controls. Typical histopathology is illustrated in Figure 4.

**Figure 4:**
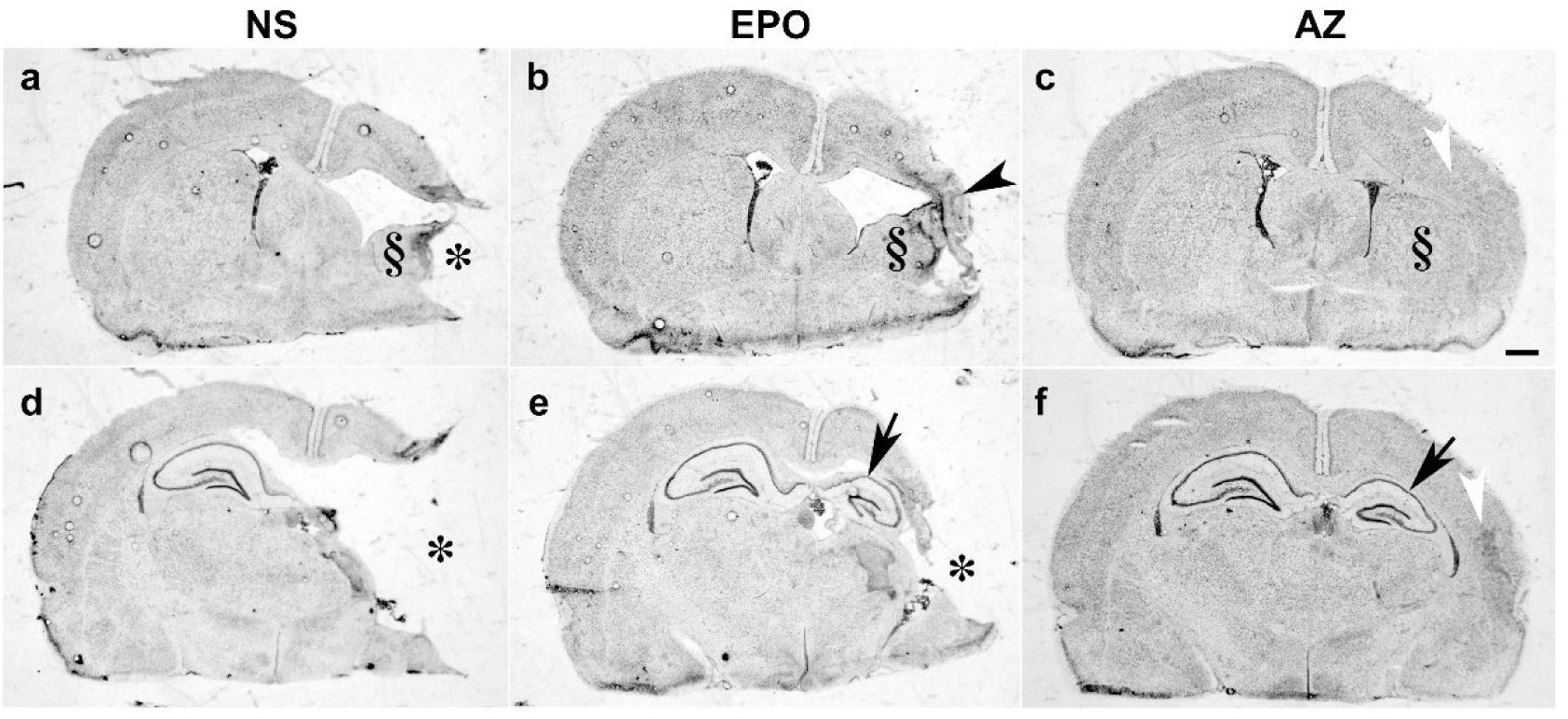
Neuropathology after Right Carotid Ligation and 90 Minutes in 8% Oxygen (Hypoxia-Ischemia): Impact of Pharmacotherapy. Animals underwent right carotid artery ligation followed by 90 min in 8% O_2_ on P7. Five-dose, once/day regimens of erythropoietin (EPO), azithromycin (AZ), or saline (NS) began 2 h post-HI, with the final dose 96h post-HI. Animals were euthanized after the completion of sensorimotor testing on P35, and brains were processed for cresyl violet staining and image analysis. Sections at the level of striatum (**a**, **b**, **c**) and dorsal hippocampus (**d**, **e**, **f**) from one representative brain from each treatment group are presented; for each brain shown, the calculated %Intact Right Hemisphere and P35 Composite Score approximated the group mean value. Moderate-severe right hemisphere damage was typical of saline-injected (NS) controls (**a**, **d**), with right cortical liquefactive infarction (*), marked striatal atrophy (§), and either marked atrophy or loss of right hippocampus. Among EPO-treated animals (**b**, **e**), right hemisphere injury was less marked than in saline controls, but cortical thinning (black arrowhead) or cystic infarction (*) were common, as was hippocampal atrophy (arrow). Striatal atrophy (§) was less marked than in controls. Among AZ-treated animals (**c**, **f**) the predominant finding was an overall relative reduction in right hemisphere volume, with right hippocampal atrophy (arrow), reduced striatal volume (§), and disruption of cortical lamination (white arrowhead) but no discernible tissue infarction. Composite Scores were 7, 22 and 25, respectively, for these representative NS, EPO and AZ-injected animals. (Scale bar 1 mm).

Second-tier outcomes that included cognitive testing were evaluated in experiments with hypothermia-treated HI, in inflammation-amplified HI without hypothermia (relevant to low-middle-income settings in which hypothermia is not safe or effective^49–51^), and hypothermia- treated inflammation-amplified HI.

#### HI plus Hypothermia

On P7 animals underwent 90 min HI and hypothermia was initiated 1h later. Five-day regimens of EPO, azithromycin or saline (16/group) began 2h after HI (i.e. 1h after initiation of hypothermia). The only deaths were in the saline group (3/16) (p = NS). After completion of sensorimotor testing on P35, animals underwent 4 days of Watermaze place navigation training, followed 24h later by a 60 sec probe trial (no platform). Sensorimotor function, % intact right hemisphere and mean Composite Score were not significantly different between the EPO and azithromycin groups; outcomes in both were significantly better than saline controls (p<0.0001 ANOVA with Tukey’s multiple comparisons test) (see Fig. 2 e). In the Watermaze probe test of memory retention, mean time to reach the former location of the platform was not significantly different between the two drug groups (Mean ±SD seconds: EPO 26.5 ±20.1; azithromycin 33.7 ±21.3).

#### Inflammation-amplified HI

Animals underwent LPS-amplified HI on P7 and received treatment with 5-day regimens of EPO, azithromycin or saline (12/group) beginning 2h after HI. Sensorimotor function was evaluated on P35; they began four days of Watermaze place navigation training on P39-40, followed 24 h later by a 60 sec probe trial, and histopathology was then evaluated. Mortality was higher (p<0.05 Fisher’s exact test) in the saline controls (4/12) than in either drug group (both 0/12). The EPO and AZ groups were similar by sensorimotor testing, hemisphere volume, and Composite Score (See Fig. 5); the AZ group had a lower mean Probe latency than the EPO group (i.e. better memory; *p<0.05 ANOVA with post-hoc Šidàk’s test). Typical features of inflammation-amplified HI histopathology in the three groups are illustrated in Figure 6; there is severe tissue damage in the right hemisphere (ipsilateral to carotid ligation) in the representative control but minimal evidence of ipsilateral tissue loss in both representative drug-treated animals. Panels **b** and **e** illustrate the greater impact of EPO treatment on severity of brain damage in LPS-amplified HI than in pure HI (see Fig. 4 **b** and **e**). Such a differential impact on neuropathology severity between models was not apparent for azithromycin.

**Figure 5:**
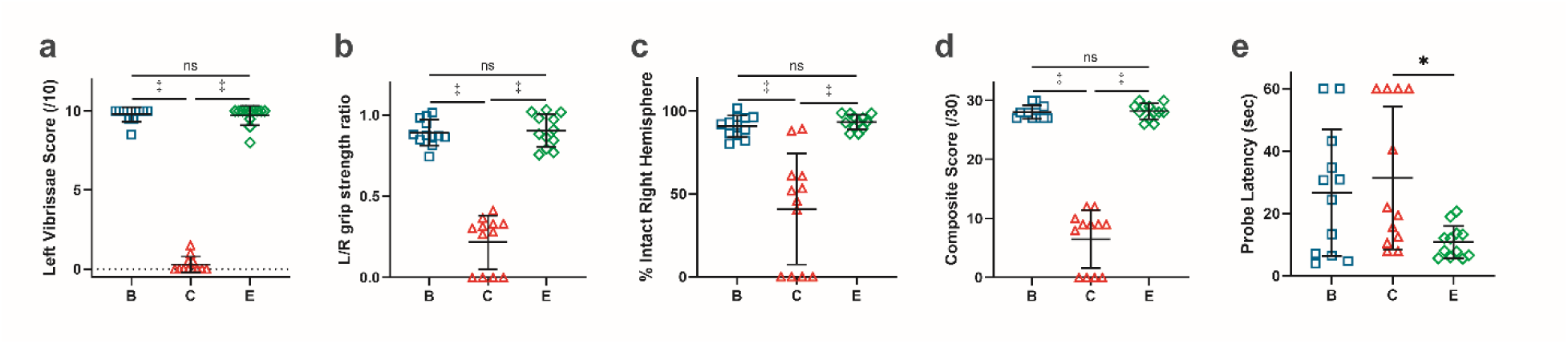
Inflammation-amplified HI; P35 Outcome: P7 rats received LPS 4h before 50 min HI. Five-Dose, once/day regimens of EPO (drug B), AZ (drug E), or saline (drug C) (12/group) began 2 h post-HI. Sensorimotor function (**a, b**) was evaluated on P35. %Intact Right hemisphere (**c**) was calculated from bilateral hemisphere volumes, and the 30-point Composite Score was calculated (**d**). Groups B and E, both superior to controls (*p<0.0001 ANOVA with Tukey’s multiple comparisons test), had similar results in sensorimotor testing, %Intact Right Hemisphere, or Composite Score (**a-d**). However, cognitive testing (Watermaze probe test) (**e**) revealed better memory in group E (azithromycin) than in group B (erythropoietin). Error bars: mean ±SD.

**Figure 6:**
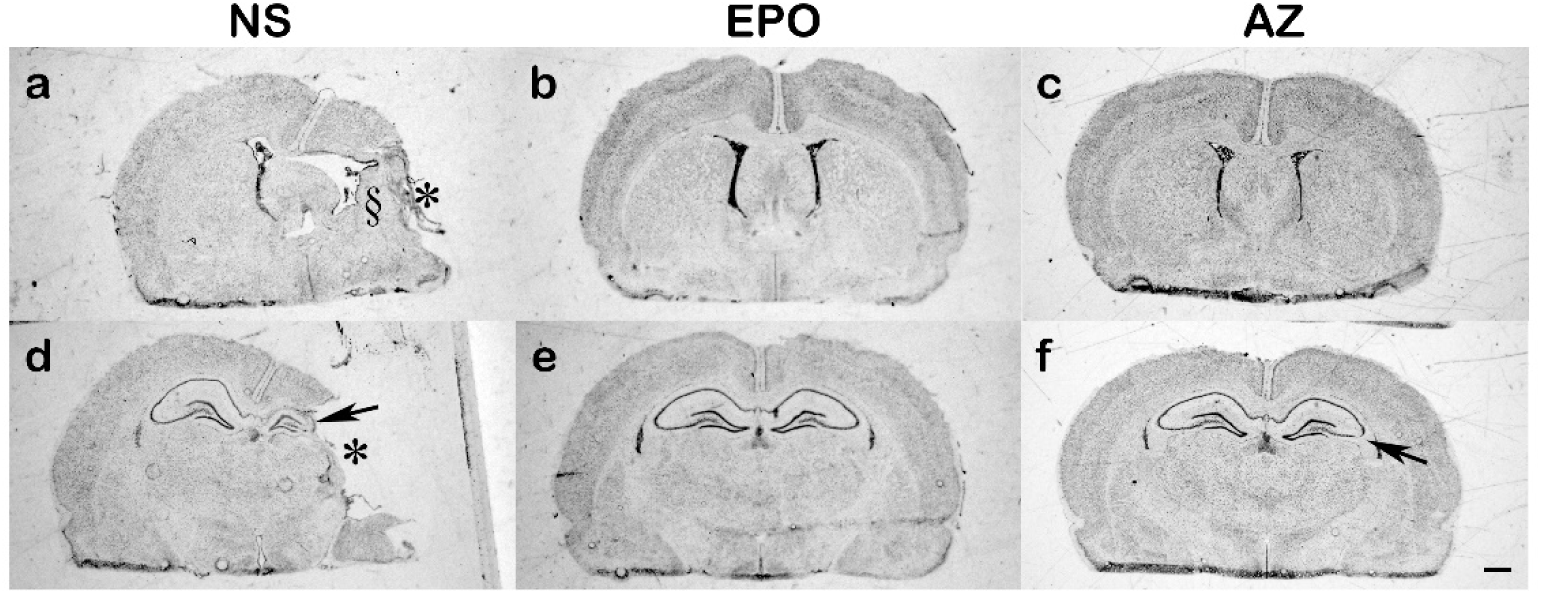
Neuropathology of Neuroprotection after LPS-Amplified Hypoxia-Ischemia: P7 rats received an i.p. LPS injection 4h before 50 min HI, in experiments summarized in Fig. 5. Five-dose, once/day regimens of erythropoietin (EPO), azithromycin (AZ), or saline (NS) began 2 h post-HI, with the final dose 96h post-HI. Animals were euthanized after the completion of Watermaze probe testing, at P43-44, and brains were processed for cresyl violet staining and image analysis. Sections from one representative brain from each treatment group are presented; for each brain the calculated %Intact Right Hemisphere approximated the group mean value shown in Fig. 5 c. Moderate-to-severe right hemisphere damage was typical of surviving saline-injected controls (**a**, **d**), with right cortical liquefactive infarction (*), right hippocampal atrophy (arrow, **d**) and left striatal infarction or atrophy (§, **a**). Among the EPO (**b**, **e**) or AZ (**c**, **f**) treated animals there was no discernible right-sided tissue infarction, with subtle relative reduction in right hemisphere volume, and in some cases, as in the AZ-treated animal shown, localized thinning of the right hippocampal pyramidal cell layer (arrow, **f**). While azithromycin treatment was associated with substantial ipsilateral tissue sparing in both pure HI and LPS-amplified HI, with EPO treatment there was marked ipsilateral tissue sparing in LPS-amplified HI rats, but less sparing of neuropathology with EPO treatment in rats that underwent HI only (compare this figure to Fig. 4). Composite scores were 9, 30 and 28, respectively, for these representative NS, EPO and AZ-injected animals. (Scale bar 1 mm)

In animals that underwent hypothermia-treated LPS-amplified HI on P7, with 5-day regimens of EPO, azithromycin or saline (8/group) beginning 2h after HI, mortality in all groups was relatively high (4/8 controls; 2/8 in each of EPO and azithromycin groups; p=NS, Fisher’s exact test). Azithromycin-treated animals had higher Composite Scores than controls (22 ±13 vs. 7 ±7 respectively, p<0.05 ANOVA with Tukey multiple comparisons test); EPO treatment benefit was borderline (mean ±SD Composite Score 21 ±13, p=0.0503 vs. control). EPO and azithromycin composite scores were similar, and Watermaze probe latency to target did not differ between the two drug groups with deaths included (mean ±SD seconds: EPO 28.8 ±26.2; azithromycin 28.7 ±26.6; deaths assigned latency = 60 sec) or deaths excluded (not shown).

## Discussion

Of the four drugs tested, in all models either azithromycin or EPO offered superior neuroprotection at the early, weaning-age outcome, with the other being the “runner-up”. Caffeine and melatonin regimens conferred modest neuroprotection in the pure HI model and the PAM-amplified HI models but were quickly eliminated by the algorithm in hypothermia-treated and LPS-amplified models. At P35-P45 azithromycin and EPO outcomes were generally similar; the only exception was that in the LPS-amplified HI model for the cognitive measure azithromycin was superior to EPO treatment.

Strengths of our approach, consistent with recent Stroke Treatment Academic Industry Roundtable (STAIR) guidelines^32^ are that we included both functional and brain damage outcomes and accounted for all deaths^25^. Also consistent with STAIR guidelines, drugs were evaluated in multiple clinically relevant models. Death is an important marker of adverse drug effects; we also evaluated serial core temperatures and weights, and blood glucose testing in inflammation-amplified HI. It was not feasible to include real-time physiological monitoring, e.g. heart and respiratory rate, oxygen saturation, or transcutaneous PCO_2_.

We mitigated the impact of known variability (within- and between-litters) in rodent HI models and the impact of investigator bias by masked block randomization within all litters. We accounted for potential sex differences by allocating both sexes to every group in all litters.

Investigators and staff performing procedures or testing remained unaware of drug-treatment identity throughout the study, until after completion of all outcome analyses. We did not preselect or exclude any animals (e.g. by evaluating early MRIs) and this approach parallels the broad range of outcome variability after HIE in clinical neonatology practice.

With a maximum litter size of 12 animals, block randomization and equal sex allocation, it was not practical to concurrently compare more than four drugs plus controls. In several injury models, our response-adaptive allocation algorithm consistently quickly eliminated drug regimens with limited potential (caffeine and melatonin), but it was more challenging to distinguish between two drugs with similar high efficacy, EPO and azithromycin. This is a reported limitation of Bayesian response-adaptive randomization; it may perform best when there is only one highly effective treatment^52^.

Azithromycin pharmacokinetics and safety in very preterm infants were reported in studies that focused on pulmonary outcomes, using doses up to 20 mg/kg/day for up to 3 days ^53–58^; none incorporated hypothermia. No neonatal safety issues were reported in several randomized controlled trials of intrapartum or neonatal azithromycin in low-middle income^59–61^ or high- income countries^62^. Safety and modest neuroprotective efficacy of azithromycin was recently demonstrated in a neonatal sheep asphyxia-resuscitation model (without hypothermia) that included motor outcomes^63^. Azithromycin could be an excellent candidate to advance to clinical trials for HIE, but some caveats must be considered. Azithromycin can cause QT interval prolongation in humans^64^; its use is associated with a risk of potentially dangerous cardiac rhythm disturbances in a few adults^65^. Its use in a randomized trial in premature infants was not associated with QT prolongation^56^. Accumulated data on QT prolongation in neonates undergoing therapeutic hypothermia was summarized in a recent report, this prolongation was asymptomatic with no severe arrhythmias and with QT normalization after re-warming^66^. Since rats have limited utility for modeling arrhythmogenic conditions^67^ azithromycin must also be tested in neonatal HI models in larger species, e.g. piglets, with thorough evaluation of arrhythmogenicity and with concurrent hypothermia.

We included EPO because these studies began before dissemination of the negative results of two large randomized trials of EPO for HIE^3,8^. The discordance between preclinical and clinical trial results is concerning, however, unlike those trials, in which EPO was initiated relatively late (mean: >16 hours after birth), we first administered all drugs at 2 hours after the end of hypoxia- ischemia. It is unclear whether the negative EPO trial results reflect delayed drug initiation or represent an inherent limitation of EPO^68^.

Caffeine improved outcome after hypoxia-ischemia in several preclinical reports ^33–35^ when dosed at 20 mg/kg caffeine citrate (or the equivalent dose of caffeine base) and in a recent comparative efficacy study in neonatal rat HI^9^, high-dose caffeine citrate (40 mg/kg, first dose pre-hypoxia) was the most effective of over two dozen drugs tested. We found caffeine citrate neuroprotection in pure HI and in PAM-amplified HI, dosed at 20 mg/kg initiated 2h after the end of hypoxia-ischemia, which could explain its lower prioritization by the algorithm. In LPS- amplified HI caffeine was rapidly dropped, suggesting lower efficacy than in pure HI or PAM- amplified HI.

Melatonin improved outcome after neonatal hypoxia-ischemia in several preclinical reports^42,43^, and it was neuroprotective when given every 12 hours starting immediately after HI in the recent report from Sabir et al. ^9^. We found melatonin neuroprotection in pure HI and PAM- amplified HI, but once-daily dosing and initial treatment at 2 h after the end of HI could explain why melatonin was de-prioritized by the algorithm. Melatonin might have fared better if dosed more frequently.

### Limitations

Translating preclinical pharmacologic neuroprotection to human infants is difficult, as is prioritization of drugs for clinical trials, as many agents are neuroprotective in preclinical models^4–6^. Repurposing FDA-approved agents eases the path to human trials. All drugs that we evaluated are FDA approved and already used in NICU patients for non-HIE indications. Our 2-hour delay from the end of the HI insult to the first drug injection acknowledges the time needed to evaluate clinical trial candidates and obtain informed consent. All dosing regimens were based on published reports of neuroprotective efficacy in rodents.

However, blood or tissue drug concentrations and pharmacokinetics were not directly evaluated. Frequency of dosing was only once/day due to staffing limitations, and we did not attempt dose optimization with dose-response or time-response studies. A potential threat to generalizability is that generic parenteral formulations of azithromycin, caffeine and erythropoietin were used, as available from our hospital pharmacy. Replicability of comparative neuroprotective efficacy between different laboratory groups, even using the same model species, may be challenging when there are differences in outcome measures or differences in generic drug preparations used. Generic drug regulations leave room for uncertainty about the extent of interchangeability among multiple generic brands^69,70^, as we recently uncovered when investigating apparent differences in neuroprotective efficacy between azithromycin preparations in our laboratory *vs*. the Sabir lab^9,24,25^.

Our use of Bayesian statistics and response-adaptive allocation was innovative, but its added value is uncertain. Adaptive designs require careful consideration and adjustment of decision rules to best allocate animals to answer questions of interest. Although our goal was to pick a “winner” it turned out that while our design quickly eliminated drugs that were neuroprotective, but not maximally so, it less readily differentiated between two treatments with fairly similar superior efficacy, erythropoietin and azithromycin. Future protocol improvements could include use of larger equal-allocation cohorts preceding initiation of response-adaptive allocation, and safeguards to avoid spending large numbers of animals in a quest to identify a single superior drug when the magnitude of difference in efficacy among remaining drugs is small and of little practical significance.

## Conclusion

Our preclinical comparative efficacy strategy tested four clinically available drugs in multiple preclinical brain injury models and consistently identified differences in efficacy among those drugs. Three key factors favoring our approach were direct within-litter comparisons, evaluation in multiple injury models and inclusion of both function and neuropathology outcomes. These results continue to support azithromycin as a candidate neuroprotective agent and support future studies in large animal neonatal cerebral hypoxia- ischemia models.

## Acknowledgements

The authors thank Jiyao Li, Mary Underwood, Krista Golden and Christopher Bidlack for masking us to drug group identities. The authors thank staff of the University of Michigan Metabolic, Physiological and Behavioral Phenotyping Core (Nathan Qi, Zhe Wu, Lauren Benson) for conducting Watermaze testing.

## Statement of Financial Support

Research reported in this publication was supported by the National Institutes of Health under award number R21 HD103869 (to J.D.E.B., W.J.M. and F.S.S.). The content is solely the responsibility of the authors and does not necessarily represent the official views of the National Institutes of Health. This research was also supported by University of Michigan Department of Pediatrics Amendt Heller Research Award and The Reiter HIE Research Fund. The Metabolic, Physiological and Behavioral Phenotyping Core is supported by the National Institutes of Health under award number U2C DK135066.

## Author Contributions

- Substantial contributions to conception and design, acquisition of data, or analysis and interpretation of data: J.D.E.B., Y.L., J.S., N.K., W.J.M. and F.S.S.
- Drafting the article or revising it critically for important intellectual content: J.D.E.B., W.J.M., F.S.S.
- Final approval of the version to be published: J.D.E.B., Y.L., J.S., N.K., W.J.M. and F.S.S.

## Competing Interests

J.D.E.B., Y.L., J.S., N.K. and F.S.S. declare no competing interests.

W.J.M. consults for Berry Consultants, Austin, TX; no consulting is related to the work presented here.

## Data Availability Statement

The datasets generated and analyzed during the current study are available from the corresponding author upon reasonable request.

## Design and Simulation Report

FACTS Core Engine for Continuous Endpoint

### 2. Introduction

#### 2.1. Background

This document describes the features of the simulated design, including the statistical models, decision rules, and simulation scenarios as input into the FACTS (Fixed and Adaptive Clinical Trial Simulator) software. A small set of operating characteristics for the simulations is also summarized.

#### 2.2. Primary Endpoint

The primary endpoint is Continuous, and is measured at 0.1 weeks. Higher response corresponds to subject improvement. The primary endpoint is a 30 point test of neurological function.

#### 2.3. Treatment Arms

The trial will enroll up to a maximum of 120 subjects, randomized among 5 arms, including a control arm. We have 4 treatment arms which we label generically by their arm index as: 𝑑 = 0 (control), 1, 2, 3, 4.

Each arm represents a different treatment regimen (drug agent or combination of drug agents).

The trial will enroll subjects in cohorts. The first cohort will consist of 24 subjects. Subsequent cohorts will each contain 12 subjects. A total of 9 cohorts may be enrolled (including the first).

### 3. Statistical Modeling

This section describes the statistical modeling used in the design. The modeling is Bayesian in nature.

#### 3.1. Final Endpoint Model

Let 𝑌_𝑖_ be the primary outcome measured as defined in the experimental protocol for the 𝑖^𝑡ℎ^ subject. We model the outcomes as

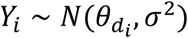

where 𝜃_𝑑_ is the mean response for arm 𝑑..

The mean response is modeled independently for each dose as:

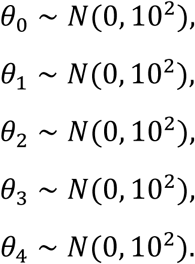

Thus, 𝜃_𝑑_ for each dose is estimated separately using only data from that dose. The error variance is modeled as:

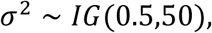

where IG(𝑎, 𝑏) is the inverse gamma distribution defined by:

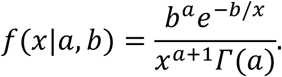

#### 3.2. Evaluation of Posterior Estimates

The Bayesian final endpoint model is fitted to the data at each update. The posterior is calculated as:

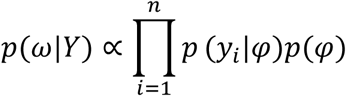

where 𝜑 is the set of parameters for the final endpoint model, 𝑝(𝜑) is the prior for those parameters, 𝑦_𝑖_ is the final response for each subject, and 𝑛𝑛 is the number of subjects. The posterior is evaluated using MCMC with individual parameters updated by Metropolis Hastings (or Gibbs sampling where possible), using only the 𝑦_𝑖_ data available at the time of the update.

#### 3.3. Quantities of Interest

We define a number of quantities that will be tracked and may be used to make decisions during the trial.

##### 3.3.1. Posterior Probabilities

For each dose, we calculate the following quantities from the posterior:

- the probability that the mean response on dose 𝑑 is greater than on control:

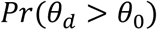

##### 3.3.2. Frequentist **𝒑**-values

We test each active dose relative to control using a one-sided two-sample t-test and calculate the 𝑝-value where missing data is ignored and the analysis is unadjusted for multiple comparisons:

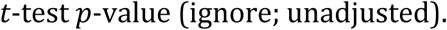

In actual study conduct, we will use 2-sided p values, and the threshold value will be adjusted for multiple comparisons.

##### 3.3.3. Target Doses

We define the following target doses:

- The maximum effective dose (𝑑_𝑚𝑎𝑥_) is the dose with the greatest treatment effect (difference from control). For each dose, we calculate the probability of being the 𝑑𝑚𝑎𝑥:

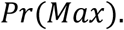

##### 3.3.4. Decision Quantities

The above quantities are computed for each treatment arm (thus making them vector quantities). To facilitate decisions in the trial, we attach a particular treatment arm to the quantity that will be used for the decision (thus reducing the vector quantity to a scalar quantity).Throughout the trial, decisions may be based on the following quantities:

- 𝑃r(𝜃_𝑑_ > 𝜃_0_) for 𝑑 = greatest 𝑃r(𝑀𝑎𝑥)
- 𝑃r(𝑀𝑎𝑥) for 𝑑 = greatest 𝑃r(𝑀𝑎𝑥)

#### 3.4. Conventions for Missing Data

At any analysis, some subjects may have missing data for the final endpoint. The missing data could result from the subject dropping out of the study, or because the subject simply has not yet reached the final visit.

For frequentist 𝑝-values, the method for handling missing data has been specified in the section Quantities of Interest. Here we describe how missing data will be handled in the Bayesian modeling.

If the subject has not yet reached the final visit (i.e. early death), the endpoint value is imputed from the estimate of the response for the subjects treatment arm (effectively contributing no information to the update of that estimate). We may conduct sensitivity analyses where we include deaths with a value of 0 on the primary endpoint.

For any subject whose final endpoint is unknown due to drop out (death), the final outcome will be multiply imputed from the Bayesian model.

### 4. Study Design

#### 4.1. Timing of Interim Analyses

Interim analyses will be conducted after every 1 cohorts complete, beginning after 1 cohorts complete. Thus, there are 8 planned interims for the trial.

#### 4.2. Response Adaptive Randomization

The first cohort of 24 subjects will be randomized in a ratio of 4:5:5:5:5. Subsequent cohorts will each enroll 12 subjects. After the first cohort, adaptive randomization will begin, with the goal of preferentially allocating subjects to the doses that appear more promising.

Within each cohort, 3 subjects will be allocated to control. For the remaining doses, we calculate:

The response adaptive allocation uses the following weights:

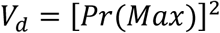

The randomization probabilities for the adaptively allocated arms will be updated at each interim. They will be weighted according to the 𝑉_𝑑_ and the weights will be renormalized to sum to 1.

To avoid assigning subjects to a dose with a minimal chance of being the best dose, any probability less than 0.1 is set to zero at that interim and the resulting probability is reallocated among the remaining doses. In this manner, a dose may be temporarily dropped but may be re-introduced if the adaptive randomization probability increases at subsequent interims.

#### 4.3. Criteria for Stopping Accrual

##### 4.3.1. Stopping for Expected Futility

For interim 1-8, the trial may stop accrual for expected futility if all of the following criteria are satisfied:

- 𝑃r(𝑀𝑎𝑥) < 0.5 for 𝑑 = greatest 𝑃r(𝑀𝑎𝑥)

This means, that if the current best dose has a lower than 50% chance of being the maximum, the trial will terminate early. This suggests that there will not be separation of the arms.

If a futility stopping rule is met at an interim analysis, then subject follow up will be discontinued, and the final evaluation criteria will be applied to the currently available data.

##### 4.3.2. Stopping for Expected Success

For interim 1-8, the trial may stop accrual for expected success if all of the following criteria are satisfied:

- 𝑃r(𝜃_𝑑_ > 𝜃_0_) > 0.99 for 𝑑 = greatest 𝑃r(𝑀𝑎𝑥)
- 𝑃r(𝑀𝑎𝑥) > 0.95 for 𝑑 = greatest 𝑃r(𝑀𝑎𝑥)

If a success stopping rule is met at an interim analysis, then a final analysis will be conducted after all currently enrolled subjects have been followed to their final endpoint.

#### 4.4. Final Evaluation Criteria

At the final analysis, the trial will be considered successful if all of the following criteria are satisfied:

- 𝑃r(𝜃_𝑑_ > 𝜃_0_) > 0.99 for 𝑑 = greatest 𝑃r(𝑀𝑎𝑥)

At the final analysis, the trial will be considered futile if any of the following criteria are satisfied:

- 𝑃r(𝜃_𝑑_ > 𝜃_0_) < 0.99 for 𝑑 = greatest 𝑃r(𝑀𝑎𝑥)

### 5. Simulation Scenarios

We evaluate the proposed design through trial simulation. We hypothesize several possible underlying truths for the mean response, as well as for trial execution variables such as accrual and dropout. For each of these scenarios, we generate data according to those truths and run through the design as specified above. We repeat this process to create multiple “virtual trials” and we track the behavior of each trial. In this section, we describe the parameters used to generate the virtual subject-level data.

#### 5.1. Virtual Subject Response Profiles

We consider 3 profiles for which subject outcomes for the final endpoint are simulated to have means as shown in Table-1 and standard deviations shown in Table-2.

**Table-1:**
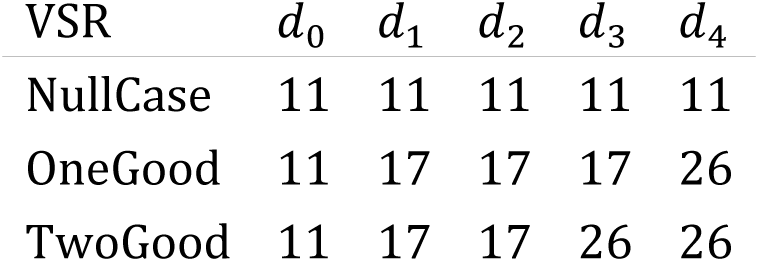
Virtual subject response means.

| VSR | $d_0$ | $d_1$ | $d_2$ | $d_3$ | $d_4$ |
| --- | --- | --- | --- | --- | --- |
| NullCase | 11 | 11 | 11 | 11 | 11 |
| OneGood | 11 | 17 | 17 | 17 | 26 |
| TwoGood | 11 | 17 | 17 | 26 | 26 |

**Table-2:** Virtual subject standard deviations.

| VSR | $d_0$ | $d_1$ | $d_2$ | $d_3$ | $d_4$ |
| --- | --- | --- | --- | --- | --- |
| NullCase | 4 | 4 | 4 | 4 | 4 |
| OneGood | 4 | 4 | 4 | 4 | 3 |
| TwoGood | 4 | 4 | 4 | 3 | 3 |

#### 5.2. Accrual Profiles

We assume that the time to accrue one cohort is 1 week.

#### 5.3. Dropout Profiles

For this study, we consider a death of an animal a dropout.

We simulate subjects dropping out of the trial with the rates (per dose and visit) shown in Table-3.

**Table-3:** Dropout Profiles.

| Profile | Week | $d_0$ | $d_1$ | $d_2$ | $d_3$ | $d_4$ |
| --- | --- | --- | --- | --- | --- | --- |
| Drop1 | 0.1 | 0.1 | 0.05 | 0.05 | 0.05 | 0.05 |

### 6. Operating Characteristics

For the scenarios described above, we simulate multiple virtual trials and track the behavior of each trial, including the final outcome of the trial, the estimated mean response, etc. The results in this section are summarized across all simulated trials for each scenario.

#### 6.1. Overall

This section gives a high-level description of the operating characteristics. Table-4 shows the following information per scenario:

- N sim: the number of simulated trials
- E[N]: the expected sample size
- Pr(success): the proportion of trials that met the final success criteria
- E[duration]: the expected duration of the trial in weeks.

**Table-4:**
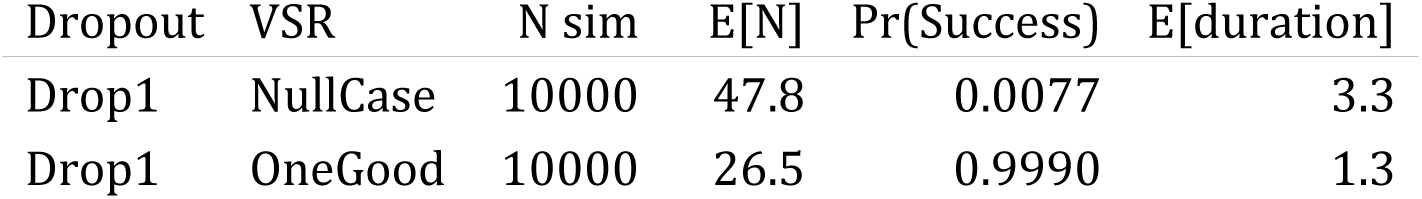
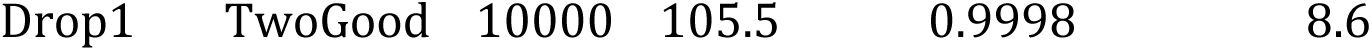
Overall Operating Characteristics.

#### 6.2. Trial Outcomes

This section summarizes the outcomes of the simulated trials. For each scenario in Table-5, the columns represent the proportion of simulated trials meeting each of the following definitions:

- Early Success (ES): stopped accrual for expected success and successful at the final analysis
- Late Success (LS): enrolled to the maximum sample size and successful at the final analysis
- Early Futility (EF): stopped accrual for futility, and met futility criteria at the final analysis
- Late Futility (LF): enrolled to the maximum sample size and met the futility criteria at the final analysis

**Table-5:** Trial Outcomes.

| Dropout | VSR | ES | LS | EF | LF |
| --- | --- | --- | --- | --- | --- |
| Drop1 | NullCase | 0.0012 | 0.0041 | 0.8574 | 0.1346 |
| Drop1 | OneGood | 0.9987 | 0.0001 | 0.0001 | 0.0000 |
| Drop1 | TwoGood | 0.2186 | 0.7701 | 0.0002 | 0.0000 |

### 7. Computational Details

This report reflects the design parameters contained within the neonatal5arm27DJan2020.facts file. The simulations were run using FACTS (Berry Consultants, LLC, Austin, TX) version 6.2.5. Table-6 shows the computational details for each scenario, including the starting date and time, the length of the MCMC chain, the random number seed, and the trial at which the simulation started. The R software package was used to summarize the simulation output and to create tables for this report.

**Table 6:**
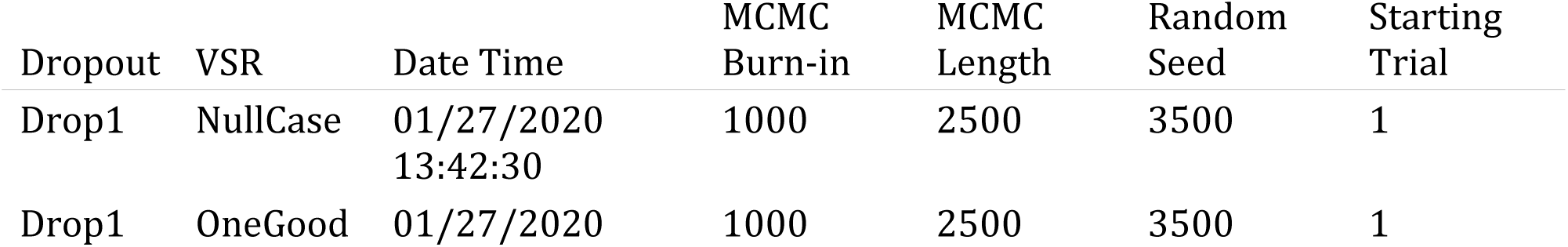
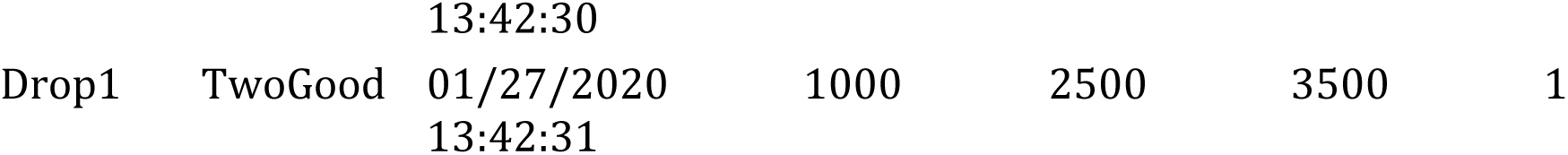
Computational Details.

**Supplemental Data Table 1.**
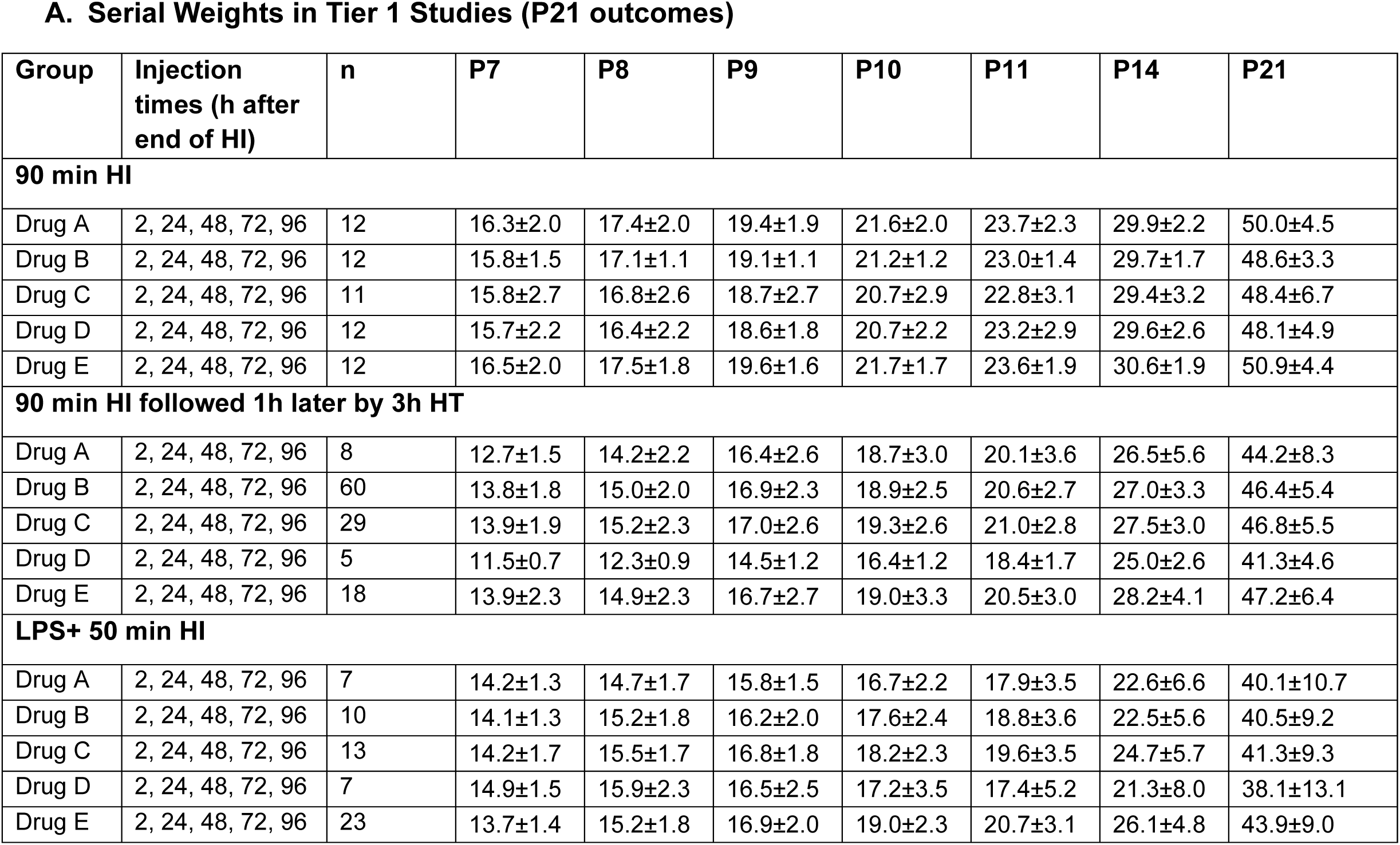
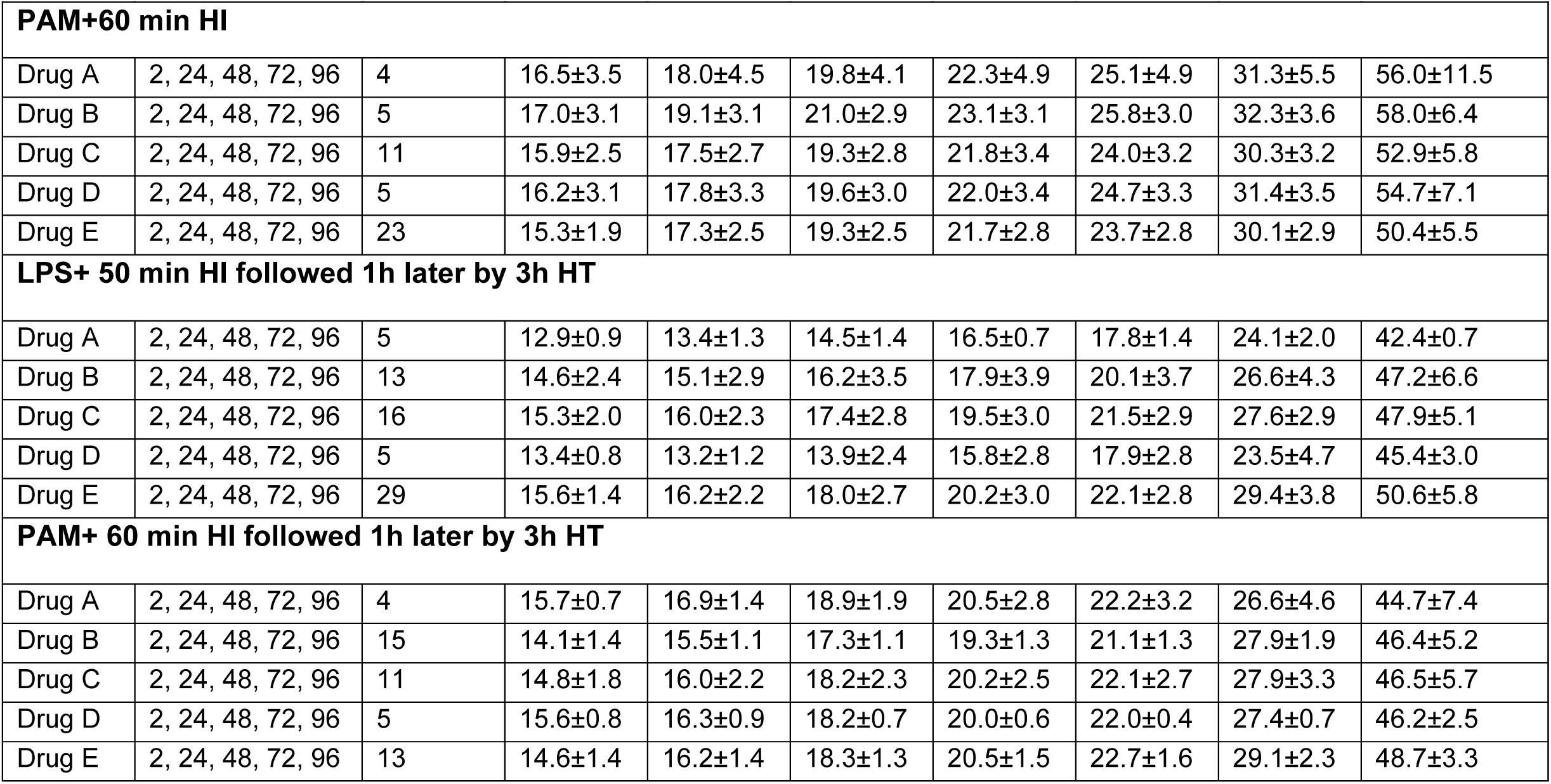
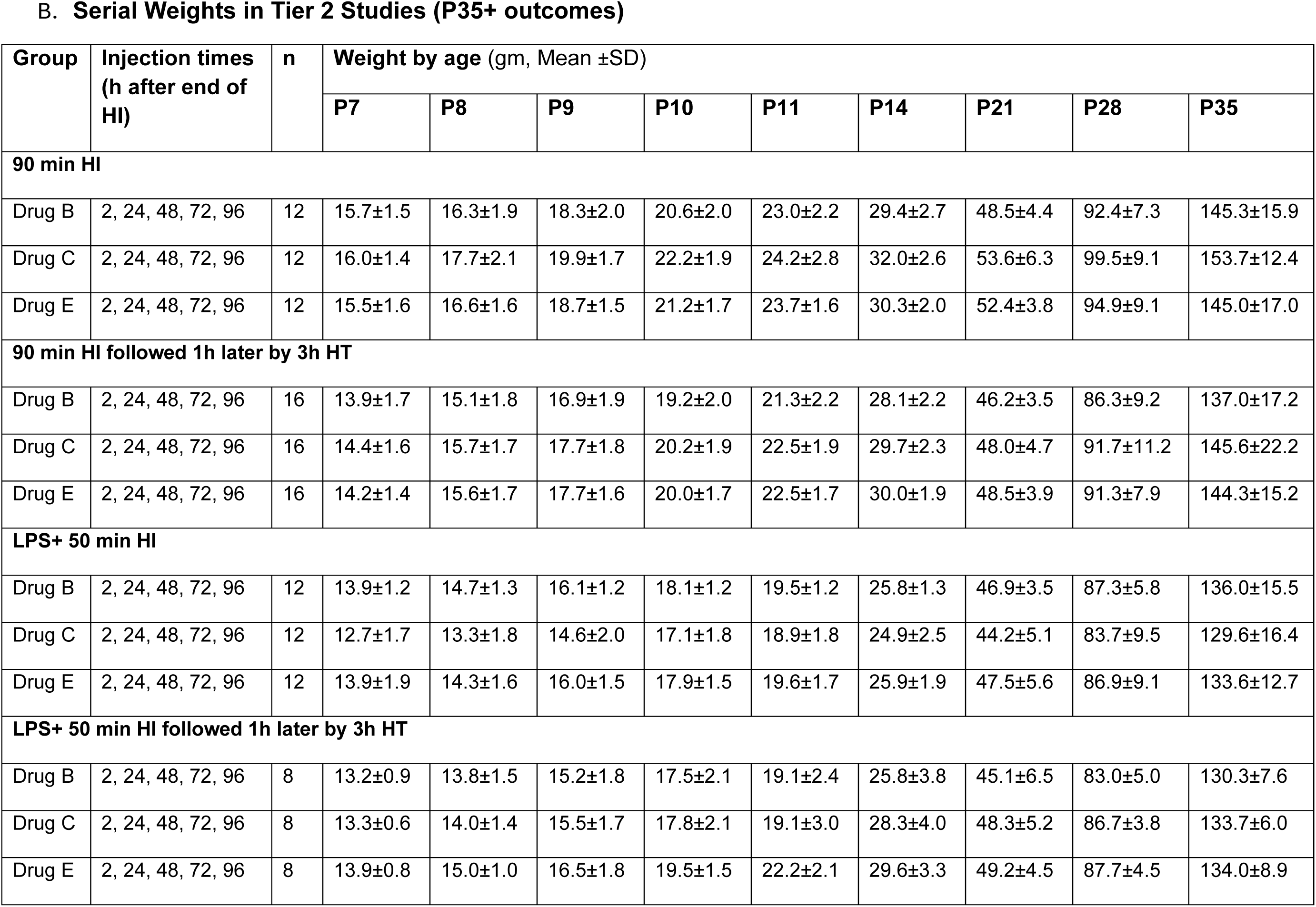

**Supplemental Data Table 2.**

| Group | Injection times<br>(h after end of<br>HI) | n | End of 8%<br>O<sub>2</sub><br>exposure | 15 min<br>after<br>hypoxia | 60 min<br>after<br>hypoxia | 120 min<br>after<br>hypoxia | 24 h<br>after<br>hypoxia | 48 h<br>after<br>hypoxia | 72 h<br>after<br>hypoxia | 96 h<br>after<br>hypoxia |
| --- | --- | --- | --- | --- | --- | --- | --- | --- | --- | --- |
| **90 min HI** |  |  |  |  |  |  |  |  |  |  |
| Drug A | 2, 24, 48, 72, 96 | 12 | 35.0±0.8 | 35.3±0.7 | 36.4±0.3 | 35.1±1.0 | 35.5±0.3 | 35.8±0.6 | 35.8±0.7 | 35.3±0.6 |
| Drug B | 2, 24, 48, 72, 96 | 12 | 34.5±0.9 | 35.5±0.9 | 36.3±0.3 | 34.9±1.3 | 35.2±0.5 | 35.7±0.5 | 35.6±0.8 | 35.4±1.1 |
| Drug C | 2, 24, 48, 72, 96 | 11 | 34.6±0.8 | 35.5±0.7 | 36.2±0.3 | 35.2±1.1 | 35.6±0.4 | 35.7±0.7 | 35.7±0.7 | 35.6±0.8 |
| Drug D | 2, 24, 48, 72, 96 | 12 | 34.7±1.0 | 35.4±0.6 | 36.5±0.4 | 35.5±0.7 | 35.1±0.6 | 35.4±0.6 | 35.6±0.7 | 35.3±0.9 |
| Drug E | 2, 24, 48, 72, 96 | 12 | 34.9±0.9 | 35.3±0.7 | 36.1±0.6 | 35.1±1.1 | 35.5±0.5 | 35.7±0.5 | 35.8±0.6 | 35.8±0.3 |
| **90 min HI followed 1h later by 3h HT** |  |  |  |  |  |  |  |  |  |  |
| Drug A | 2, 24, 48, 72, 96 | 8 | 35.2±0.7 | 35.2±0.4 | 35.9±0.3 | 30.5±0.5 | 35.4±0.6 | 35.2±0.6 | 35.1±0.6 | 34.6±1.3 |
| Drug B | 2, 24, 48, 72, 96 | 60 | 34.9±0.8 | 35.6±0.6 | 36.3±0.4 | 31.2±0.7 | 35.4±0.7 | 35.4±0.7 | 35.3±0.8 | 35.2±0.6 |
| Drug C | 2, 24, 48, 72, 96 | 29 | 35.0±0.7 | 35.4±0.6 | 36.3±0.5 | 31.1±0.7 | 35.5±0.6 | 35.3±0.7 | 35.3±0.9 | 35.5±0.6 |
| Drug D | 2, 24, 48, 72, 96 | 5 | 34.3±0.6 | 35.3±0.6 | 36.2±0.3 | 30.6±0.5 | 35.2±0.4 | 35.1±0.5 | 35.2±0.6 | 34.5±1.2 |
| Drug E | 2, 24, 48, 72, 96 | 18 | 34.7±0.6 | 35.4±0.7 | 36.2±0.5 | 30.8±0.7 | 35.4±0.5 | 35.3±0.6 | 35.1±0.5 | 35.2±0.7 |
| **LPS+ 50 min HI** |  |  |  |  |  |  |  |  |  |  |
| Drug A | 2, 24, 48, 72, 96 | 7 | 35.0±1.0 | 35.1±0.6 | 36.4±0.6 | 35.5±0.8 | 35.5±0.6 | 35.3±1.2 | 34.0±2.6 | 36.0±0.8 |
| Drug B | 2, 24, 48, 72, 96 | 10 | 35.6±0.6 | 35.4±0.6 | 36.3±0.5 | 35.0±1.0 | 35.5±0.6 | 35.5±0.5 | 35.2±1.3 | 36.1±0.7 |
| Drug C | 2, 24, 48, 72, 96 | 13 | 35.3±0.6 | 35.5±0.6 | 36.5±0.3 | 35.5±0.7 | 35.5±0.4 | 35.5±0.5 | 35.0±1.7 | 35.7±0.7 |
| Drug D | 2, 24, 48, 72, 96 | 7 | 35.6±0.6 | 35.6±0.5 | 36.2±0.6 | 35.0±1.9 | 35.3±0.9 | 35.5±0.5 | 32.9±2.9 | 35.8±0.4 |
| Drug E | 2, 24, 48, 72, 96 | 23 | 35.3±0.6 | 35.3±0.5 | 36.0±0.5 | 35.0±1.2 | 35.4±0.5 | 35.5±0.6 | 35.5±1.8 | 35.7±0.5 |
| **PAM+60 min HI** |  |  |  |  |  |  |  |  |  |  |
| Drug A | 2, 24, 48, 72, 96 | 4 | 36.0±0.6 | 35.8±0.3 | 36.8±0.3 | 36.0±0.7 | 36.1±0.3 | 35.3±0.3 | 36.0±0.4 | 36.4±0.3 |
| Drug B | 2, 24, 48, 72, 96 | 5 | 35.7±0.3 | 36.3±0.3 | 36.5±0.4 | 35.7±0.6 | 35.9±0.4 | 35.3±0.6 | 35.6±0.4 | 36.2±0.4 |
| Drug C | 2, 24, 48, 72, 96 | 11 | 34.9±0.9 | 35.6±0.6 | 36.4±0.5 | 35.8±0.6 | 36.1±0.3 | 35.6±0.7 | 35.7±0.3 | 36.0±0.4 |
| Drug D | 2, 24, 48, 72, 96 | 5 | 35.4±0.5 | 36.0±0.5 | 36.7±0.4 | 35.9±0.7 | 35.6±0.4 | 35.0±0.6 | 35.8±0.6 | 36.2±0.4 |
| Drug E | 2, 24, 48, 72, 96 | 23 | 35.6±0.4 | 35.7±0.4 | 36.4±0.4 | 35.5±0.6 | 35.8±0.4 | 35.6±0.8 | 35.6±0.5 | 35.8±0.4 |
| **LPS+ 50 min HI followed 1h later by 3h HT** |  |  |  |  |  |  |  |  |  |  |
| Drug A | 2, 24, 48, 72, 96 | 5 | 35.4±0.4 | 35±0.6 | 36.2±0.7 | 30.2±0.3 | 35±0.9 | 34.6±0.5 | 35.5±0.9 | 36±0.7 |
| Drug B | 2, 24, 48, 72, 96 | 13 | 35.3±1.1 | 35.8±0.8 | 36.3±0.4 | 30.9±0.8 | 35.5±0.9 | 35.6±0.9 | 35.7±1.0 | 36.0±0.6 |
| Drug C | 2, 24, 48, 72, 96 | 16 | 35.7±0.8 | 35.8±0.7 | 36.7±0.5 | 31.4±0.7 | 35.7±0.7 | 35.5±0.9 | 36.1±0.8 | 35.8±1.4 |
| Drug D | 2, 24, 48, 72, 96 | 5 | 34.7±0.7 | 35.3±0.3 | 36.4±0.2 | 30.2±0.4 | 34.8±0.9 | 34.6±0.6 | 34.9±0.5 | 35.5±0.8 |
| Drug E | 2, 24, 48, 72, 96 | 29 | 35.9±0.7 | 36.0±0.7 | 36.8±0.4 | 31.6±0.8 | 35.8±0.6 | 35.5±1.0 | 36.2±0.6 | 36.2±0.5 |
| **PAM+ 60 min HI followed 1h later by 3h HT** |  |  |  |  |  |  |  |  |  |  |
| Drug A | 2, 24, 48, 72, 96 | 4 | 35.9±1.1 | 35.9±0.5 | 37.5±0.0 | 32.3±0.6 | 36.6±0.3 | 36.9±0.5 | 37.5±0.0 | 37.0±0.0 |
| Drug B | 2, 24, 48, 72, 96 | 15 | 36.5±0.6 | 36.4±0.5 | 37.1±0.5 | 31.9±0.8 | 36.2±0.7 | 36.3±0.7 | 36.5±0.6 | 36.9±0.3 |
| Drug C | 2, 24, 48, 72, 96 | 11 | 36.2±1.0 | 36.4±0.8 | 37.2±0.4 | 32.3±0.5 | 36.3±0.6 | 36.4±0.7 | 36.9±0.7 | 36.8±0.5 |
| Drug D | 2, 24, 48, 72, 96 | 5 | 36.0±0.6 | 35.9±0.7 | 37.4±0.2 | 32.3±0.4 | 36.7±0.4 | 36.9±0.8 | 37.5±0.0 | 36.7±0.3 |
| Drug E | 2, 24, 48, 72, 96 | 13 | 36.7±0.7 | 36.4±0.5 | 37.1±0.3 | 32.2±0.6 | 36.2±0.5 | 36.3±0.7 | 36.8±0.7 | 36.8±0.2 |

B. Serial Temperature Measurements, Tier 2 experiments (P35+ Outcomes)
| Group | Injection times<br>(h after end of<br>HI) | n | End of 8%<br>O<sub>2</sub><br>exposure | 15 min<br>after<br>hypoxia | 60 min<br>after<br>hypoxia | 120 min<br>after<br>hypoxia | 24 h<br>after<br>hypoxia | 48 h<br>after<br>hypoxia | 72 h<br>after<br>hypoxia | 96 h<br>after<br>hypoxia |
| --- | --- | --- | --- | --- | --- | --- | --- | --- | --- | --- |
| **90 min HI** |  |  |  |  |  |  |  |  |  |  |
| Drug B | 2, 24, 48, 72, 96 | 12 | 35.6±0.4 | 36.2±0.4 | 36.7±0.2 | 35.6±0.9 | 35.5±0.5 | 35.8±0.5 | 36.2±0.3 | 36.3±0.5 |
| Drug C | 2, 24, 48, 72, 96 | 12 | 35.6±0.6 | 35.9±0.6 | 36.5±0.3 | 35.4±0.6 | 35.6±0.2 | 35.9±0.5 | 36.4±0.3 | 36.1±0.4 |
| Drug E | 2, 24, 48, 72, 96 | 12 | 35.6±0.9 | 35.9±0.4 | 36.7±0.3 | 35.2±0.7 | 35.5±0.7 | 35.6±0.4 | 36.3±0.3 | 35.9±0.6 |
| **90 min HI followed 1h later by 3h HT** |  |  |  |  |  |  |  |  |  |  |
| Drug B | 2, 24, 48, 72, 96 | 16 | 35.4±0.4 | 35.6±0.6 | 36.5±0.4 | 31.3±0.6 | 34.8±0.9 | 34.7±0.5 | 35.3±0.6 | 35.5±0.6 |
| Drug C | 2, 24, 48, 72, 96 | 16 | 35.3±0.5 | 35.8±0.4 | 36.5±0.3 | 31.3±0.7 | 34.9±0.6 | 34.8±0.5 | 35.5±0.5 | 35.5±0.7 |
| Drug E | 2, 24, 48, 72, 96 | 16 | 35.2±0.5 | 35.8±0.4 | 36.7±0.3 | 31.3±0.7 | 34.8±0.8 | 34.7±0.5 | 35.4±0.5 | 35.5±0.7 |
| **LPS+ 50 min HI** |  |  |  |  |  |  |  |  |  |  |
| Drug B | 2, 24, 48, 72, 96 | 12 | 35.4±0.6 | 35.6±0.7 | 36.4±0.4 | 35.0±1.3 | 35.4±0.8 | 35.4±0.3 | 35.3±0.4 | 35.8±0.5 |
| Drug C | 2, 24, 48, 72, 96 | 12 | 35.4±0.5 | 35.6±0.8 | 36.2±0.4 | 34.5±1.7 | 35.3±0.7 | 34.9±1.2 | 35.4±0.5 | 35.9±0.7 |
| Drug E | 2, 24, 48, 72, 96 | 12 | 35.5±0.9 | 35.7±0.7 | 36.6±0.5 | 34.2±1.8 | 35.3±0.7 | 35.5±0.3 | 35.3±0.9 | 36.1±0.6 |
| **LPS+ 50 min HI followed 1h later by 3h HT** |  |  |  |  |  |  |  |  |  |  |
| Drug B | 2, 24, 48, 72, 96 | 8 | 35.8±0.5 | 36.0±0.4 | 36.5±0.0 | 30.4±0.2 | 35.1±0.5 | 34.8±1.4 | 35.0±0.6 | 35.1±0.8 |
| Drug C | 2, 24, 48, 72, 96 | 8 | 35.8±0.6 | 36.0±0.5 | 36.4±0.3 | 30.3±0.3 | 35.2±0.9 | 35.3±0.3 | 35.0±0.5 | 35.3±1.0 |
| Drug E | 2, 24, 48, 72, 96 | 8 | 35.9±0.4 | 36.1±0.6 | 36.4±0.2 | 30.8±0.5 | 35.3±0.5 | 35.5±0.0 | 35.2±0.4 | 35.8±0.8 |

